# Genetic memory devices to detect specialized metabolites in plant and soil microbiomes

**DOI:** 10.1101/2025.01.30.635401

**Authors:** Morten L. Hansen, Kristoffer I.T. Kordatos, Jens-Jakob K. Nørgaard, Johan P.B. Jørgensen, Mikael L. Strube, Morten D. Schostag, Lei Yang, Lars Jelsbak

**Affiliations:** Department of Biotechnology and Biomedicine, Technical University of Denmark, Søltofts Plads bldg. 221, DK-2800 Kgs Lyngby, Denmark; Novo Nordisk Foundation Center for Biosustainability, Technical University of Denmark, Søltofts Plads bldg. 221, DK-2800 Kgs Lyngby, Denmark

**Keywords:** Biosensors, DAPG, pyoluteorin, tetracycline, salicylate, naringenin

## Abstract

Root-associated microbiomes significantly influence plant growth and resilience through intricate chemical dialogues mediated by plant- and microbe-derived specialized metabolites. These metabolites play pivotal roles in shaping the assembly, dynamics, and ecological functions of soil microbiomes. Despite advances in *in vitro* and DNA sequencing studies, a comprehensive understanding of *in situ* chemical signaling within plant and soil microbiomes remains elusive due to experimental constraints. To address this gap, we developed and tuned a set of five whole-cell biosensors in *Escherichia coli* for spatiotemporal, non-disruptive detection of biologically relevant specialized metabolites, including 2,4-diacetylphloroglucinol, pyoluteorin, tetracycline, salicylic acid, and naringenin. Four of these biosensors were successfully adapted to the soil-compatible *Pseudomonas putida* KT2440 Δall-Φ strain. Additionally, the four sensors were shown to respond to their cognate ligand in a non-sterile soil extract medium containing the diverse microbiome found in soil. By employing genetic memory devices with DNA barcodes for readouts, our approach provides a scalable platform for sensing additional specialized metabolites in the future. This work demonstrates the potential of real-time biosensor technologies to unravel the complex chemical interactions driving soil microbiome ecology, with implications for sustainable agricultural practices.

## Introduction

Root-associated microbiomes are well-established as beneficial contributors to plant growth and resilience against biotic and abiotic stressors, with plant- and microbe-derived secondary metabolites serving as key mediators of the molecular dialogue between microorganisms and plants in the rhizosphere, thereby playing crucial roles in maintaining plant health (1–3). Understanding the roles of these secreted metabolites in mediating the assembly, dynamics, and functional behaviors of soil and root-associated microbiomes is not only fundamental to unraveling the ecological processes governing soil microbiomes but also hold significant potential for developing innovative strategies to harness native microbiomes for sustainable crop production (4, 5). However, our current understanding of chemical signaling within soil ecology remains constrained by the inherent challenges of conducting experimental analyses *in situ*. Various experimental approaches have been employed to address this gap. For instance, *in vitro* studies have elucidated numerous instances of chemical signaling among plants and microbes (6–9); however, the ecological relevance of these findings to plant and soil microbiomes under natural conditions often remains ambiguous. Similarly, DNA sequencing of soil microbial communities has uncovered interaction patterns (10, 11), but the specific molecular signals underpinning these interactions frequently remain unidentified. Notably, direct causal links between plant- and microbe-produced signals and the functional and ecological dynamics of soil microbiomes have been established only in a limited number of cases (12, 13). Currently, no technology exists to enable non-disruptive *in situ* spatiotemporal analysis and discovery of microbial signaling within plant and soil microbiomes. Here we aim to develop such a technology based on real-time whole-cell biosensors and demonstrate its potential in enhancing our understanding of soil microbiome ecology and function.

Real-time biosensors offer a promising alternative for longitudinal studies of microbial communities by enabling continuous monitoring of bioavailable analytes within the community. Unlike traditional analytical methods, biosensors utilize living organisms to report on analyte presence and concentration over time. Conventional biosensors often rely on visual outputs for reporting (14–16). However, this approach necessitates optical access, which is a limitation in natural environments. Biosensors, capable of recording analyte presence without continuous optical access, could overcome this challenge (17). One example of this involves the use of site-specific recombinases, such as serine integrases. Integrases originate from bacteriophages, where they catalyze the insertion of phage DNA into host genomes by recombination between the *attP* (phage) and *attB* (bacterial host) attachment sites (17, 18). By placing both attachment sites on the same piece of DNA, the integrase-mediated inversion serves as genetic memory, since the recombination event catalyzed by a serine integrase is irreversible in the absence of a recombination directionality factor (19). This genetic memory can be accessed later with sequencing, which can be further multiplexed with amplicon sequencing allowing simultaneous parallel investigation across different conditions, soils and time points. By connecting the expression of the integrase to a transcription factor-based sensor, cellular memory can be used to infer exposure to a particular bioavailable chemical signal recognized by the specific transcription factor. This is useful if direct readout of a reporter from a microenvironment is not possible. This technology was previously employed to record the presence of antibiotics (20) and biomarkers of inflammatory bowel disease (21) in the mouse gut. Building on this foundation, we propose to develop genetic memory devices tailored to investigate the chemical signaling processes among plants and soil microbes, with a particular focus on specialized metabolites that mediate communication and ecological functions.

The specialized metabolites 2,4-diacetylphloroglucinol (DAPG), pyoluteorin, and tetracycline represent antimicrobial compounds relevant to biocontrol and the competitive nature of rhizosphere residents. DAPG and pyoluteorin, primarily produced by *Pseudomonas* species, have been shown to significantly influence the assembly and composition of root-associated microbiomes (3). These compounds function as antimicrobial agents, selectively inhibiting or promoting the growth of specific microbial taxa within the rhizosphere (8, 22). Tetracycline, produced by certain species of *Streptomyces*, is part of the natural arsenal of secondary metabolites produced by these soil-dwelling bacteria, which they use to inhibit the growth of competing microorganisms in their environment, thereby gaining a competitive advantage in nutrient-limited soil ecosystems (16, 23). Additionally, plants are also known to secrete small molecules, which serve as signals between plants and their associated root microbiomes, such as salicylic acid and flavonoids, including naringenin. Salicylic acid is a key metabolite involved in plant induced systemic resistance, where it also serves as a root-to-root signal between adjacent plants in response to pathogen invasion (24–26). Flavonoids, including naringenin, are a group of specialized metabolites secreted by legumes, which act as chemoattractants and inducers of nitrogen fixation in endosymbiotic *Rhizobium* species (13, 27, 28). Investigating the spatiotemporal dynamics and bioavailability of these secreted specialized metabolites *in situ* could further enhance our understanding of the intricate chemical dialogue and warfare occurring between plants and their root-associated microbiomes.

In this study, we developed a catalogue of five orthogonal biosensors using genetic memory as output to allow for non-disruptive *in situ* spatiotemporal detection of biologically relevant specialized metabolites found in soil and rhizosphere environments. The biosensors were tuned and optimized in *E. coli* and shown to respond to nano- to micromolar concentrations of the five metabolites: DAPG, pyoluteorin, tetracycline, salicylic acid, and naringenin. The sensory and reporter units of four of the biosensors, except the naringenin sensor, were chromosomally integrated into the root-colonizing *P. putida* KT2440 Δall-Φ to enable long duration *in situ* experiments without requiring antibiotic selection. Lastly, we demonstrated that the four sensors, after a final optimization step, functioned in a non-sterile soil extract medium containing the diverse microbiome found in soil.

## Results and discussion

### Engineering of a two-plasmid genetic memory device for detection of specialized metabolites

In order to detect specialized metabolites in natural environments, including soil and rhizosphere, we engineered a two-plasmid system based on Yang et al. (29) utilizing irreversible integrase-mediated DNA recombination (genetic memory) as output. The device relies on transcription factor-based recognition of its cognate inducing ligand. Here, we initially built a system with the PhlF repressor optimized by Meyer et al. (30) for detection of 2,4-diacetylphloroglucinol (DAPG) as trial system. In the absence of DAPG, the constitutively expressed PhlF-repressor binds to the operator located within the **P**_phlF_ promoter, thereby repressing transcription of *integrase12*, leaving the genetic conformation of *attB*/*attP* on pRep12 intact with the *gfp* gene in the wrong orientation for transcription (Fig. 1A, *top*). In the presence of DAPG, the repression on **P**_phlF_ by PhlF is relieved, leading to the expression of Integrase12, which in turn catalyzes irreversible recombination of *attB*/*attP* to *attL*/*attR*, which reorients the *gfp* gene to be constitutively transcribed from **P**_J23119_ (Fig. 1A, *bottom*). Biosensor cells that have undergone Integrase12-mediated recombination can thus be selected with flow cytometry first based on their red fluorescence indicating presence of pRep12 and subsequently their green fluorescence indicating successful integrase-mediated recombination.

**Figure 1.**
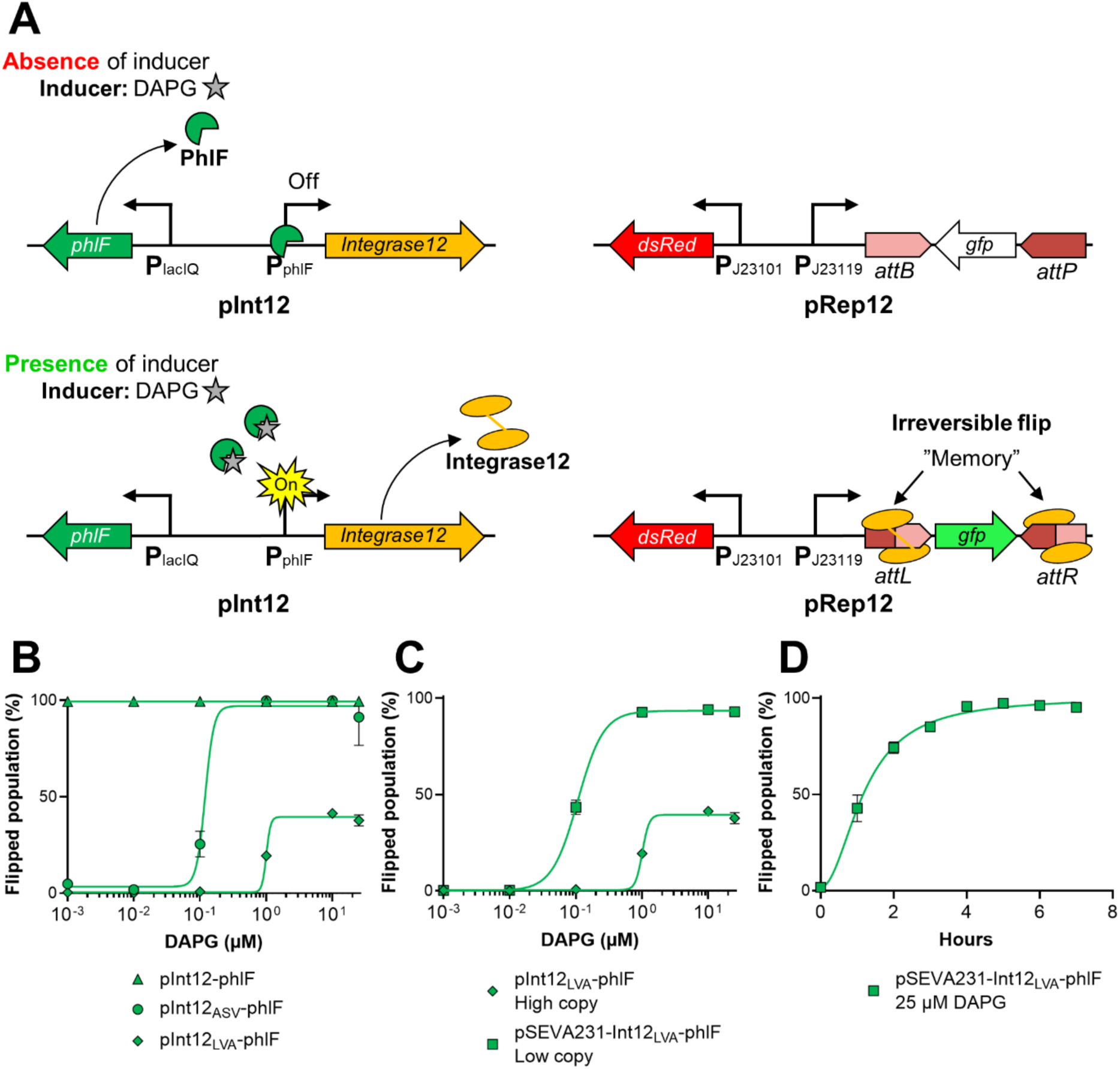
Design and optimization of a genetic memory device. **A)** The two-plasmid system comprising the genetic memory devices. *Top* In the absence of inducer, transcription of *integrase12* is off, and the biosensor cells containing both pInt12 and pRep12 emit exclusively red fluorescent light, due to expression of *dsRed* from the constitutive P_J23101_ promoter. *Bo2om* In the presence of inducer, repression of *integrase12* is relieved, leading to expression of the protein. Integrase 12 in turn catalyzes an irreversible reorientation of *a2B* and *a2P*, which flips the DNA sequence between the recognition sites 180°, resulting in biosensor cells emitting both red and green fluorescence constitutively. **B)** Response curves comparing the addition of a protein degradation tag to *integrase12* to remove the leakiness of the system in the absence of inducer. **C)** Response curves comparing the reduction of copy numbers of the plasmid containing the integrase system to match that of pRep12 increased the dynamic range, improved sensitivity, while still preventing leakiness. **D)** Response curve of the PhlF system over time in the presence of 25 µM DAPG. The left-most data point in B and C represents the recombination in the absence of inducer. Data points represent mean values of three biological replicates.

However, due to the inherent irreversible nature of recombination mediated by serine site-specific integrases, the expression of Integrase12 must be tightly controlled to avoid leakiness. Here, we opted for post-translational control by altering the stability of Integrase12 with the addition of a *ssrA*-like peptide-tag to the C-terminal end. Addition of the *ssrA*-tag directs protein degradation by the cytoplasmic ClpXP protease in *E. coli* (31). We tested two different versions of the *ssrA*-tag, where the last three amino acids were changed from the consensus sequence, AANDENYA<u>LAA</u>, to represent a weak (ASV) and a strong (LVA) degradation rate based on Andersen et al. (32). We observed that the LVA tag was required to avoid leaky behavior in the absence of DAPG, although resulting in reduced dynamic range (Fig. 1B).

Next, we sought to equalize the copy numbers of the integrase and reporter plasmids to mimic the outcome of chromosomally integrating both the sensory and reporting part, which was the desired end-goal and a necessity to obtain whole-cell biosensors that reliably propagate in natural environments without plasmid selection. The pInt12 background from Yang et al. (29) contains a ColE1 origin of replication, which results in semi-high plasmid copy numbers (30-40 copies) in *E. coli*, whereas the p15a origin of replication on pRep12 results in low copy numbers (6-10 copies) in *E. coli* (33). Thus, we integrated the sensory part of the system on pSEVA231, which carries the BBR1 origin of replication (4-6 copies in *E. coli* (33)), thereby equalizing the copy numbers of the sensory and the reporting system. This change ultimately resulted in an improved dynamic range, where the entire bacterial population responded to the presence of DAPG, as well as a 10-fold increased sensitivity (Fig. 1C). This improvement could be a result of reducing the overall intracellular levels of transcription factor, thereby requiring a lower concentration of DAPG to relieve repression, which has also been observed previously (34). Lastly, we determined the efficiency of the low-copy Integrase12-mediated recombination over time in a nutrient-rich environment. In the presence of DAPG at a concentration capable of inducing an entire population, cells respond immediately, and full population-wide induction was reached within 4 hours (Fig. 1D).

### Broadening the catalogue of memory devices detecting relevant metabolites

To detect a larger repertoire of biologically relevant metabolites, four additional biosensors were constructed to respond to pyoluteorin, tetracycline, salicylic acid, and naringenin. The pyoluteorin sensor is based on the PltZ repressor from *Pseudomonas protegens* DTU9.1, which naturally represses the transcription of the genetic operon responsible for export of pyoluteorin (35). The *pltZ* gene was codon optimized for *E. coli* and inserted in the pSEVA231-Int12_LVA_ background along with a rationally designed promoter (**P**_pltZ_) containing the PltZ-operator located between the −35 and −10 regions (Supplementary Table 5). The strength of the ribosome binding site (RBS) upstream of *pltZ* as measured by the relative translation rate was decreased by using the Salis lab prediction tool (36) (Fig. S1A), resulting in the optimal pSEVA231-Int12_LVA_-pltZ_RBS0.05x_ responding to nanomolar concentrations of pyoluteorin (Fig. 2B).

**Figure 2.**
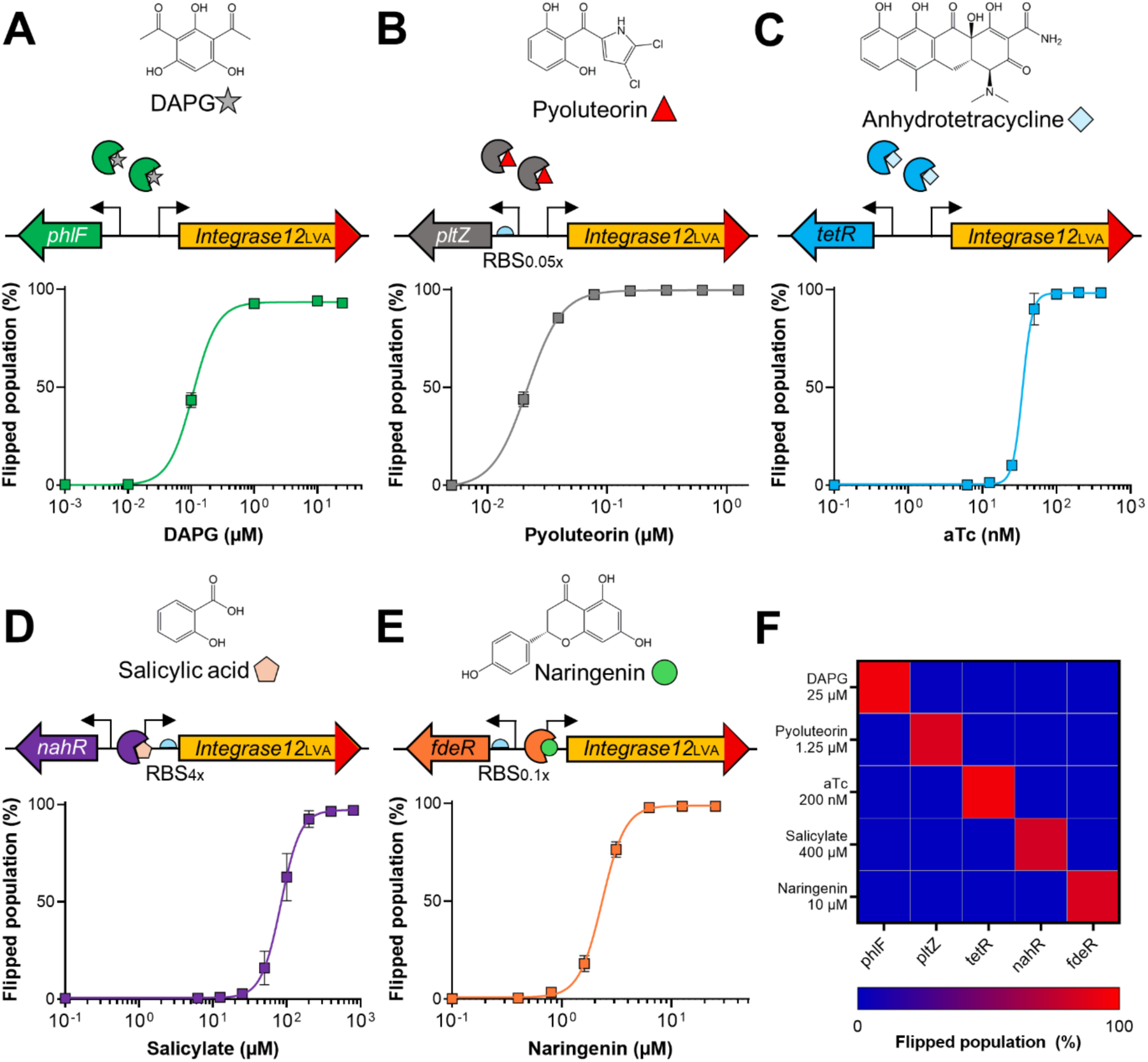
Broadening the catalogue of genetic memory devices. All biosensors contain pRep12 and a unique integrase system for each metabolite. The left-most data point represents the recombination in the absence of inducer. The red part of the *integrase12* gene indicates the presence of the C-terminal LVA-tag. **A)** Response curve of the biosensor containing pSEVA231-Int12_LVA_-phlF responding to DAPG. **B)** Response curve of the biosensor containing pSEVA231-Int12_LVA_-pltZ_RBS0.05x_ responding to pyoluteorin. **C)** Response curve of the biosensor containing pSEVA231-Int12_LVA_-tetR responding to anhydrotetracycline. **D)** Response curve of the biosensor containing pSEVA231-Int12_LVA,RBS4x_-nahR responding to salicylate. **E)** Response curve of the biosensor containing pSEVA231-Int12_LVA_-fdeR_RBS0.1x_ responding to naringenin. **F).** Orthogonality matrix for each metabolite and their respective biosensor. Data represents the average of three biological replicates.

To engineer the tetracycline and salicylic acid sensors, we integrated the TetR and NahR systems from Meyer et al. (30) into our pSEVA231-Int12_LVA_ background. For the tetracycline sensor we utilized the less toxic version, anhydrotetracycline (aTc), as the inducer. This sensor also responded to nanomolar concentrations of its inducer (Fig. 2C). The salicylic acid sensor was further tuned by optimizing the RBS upstream of Int12 to obtain a greater dynamic range (Fig. S1B), resulting in a sensor that responded to micromolar concentrations of salicylate (Fig. 2D). The naringenin sensor was based on the FdeR activator from *Herbaspirillum seropedicae* SmR1 (37). The *fdeR* gene was codon optimized for *E. coli* and integrated in the pSEVA231-Int12_LVA_ background along with the 275 bp region containing the natural **P**_fdeR_ promoter and the FdeR binding site. The RBS upstream of *fdeR* was further optimized (Fig. S1C), resulting in the optimal pSEVA231-Int12_LVA_-fdeR_RBS0.1x_ responding to low micromolar concentrations of naringenin (Fig. 2E). Lastly, we investigated the orthogonality of the five biosensors and their cognate inducers. As displayed in Fig. 2F there was no cross-talk between the specific sensors and unspecific inducers, supporting the potential application of these biosensors in a cocktail capable of accurately detecting multiple metabolites simultaneously.

### The biosensors detect live-produced metabolites and function in a soil-like environment

Next, we sought to investigate the functionality of the *E. coli* biosensors by assaying their ability to sense and respond to live-produced metabolites. We cultivated *P. protegens* DTU9.1 wildtype, as well as mutants deficient in production of either DAPG (Δ*phlACB*), pyoluteorin (Δ*pltA*), or both metabolites on agar surfaces. The DAPG and pyoluteorin biosensors both containing pRep12 and either pSEVA231-Int12_LVA_-phlF or pSEVA231-Int12_LVA_-pltZ_RBS0.05x_ were subsequently inoculated adjacent to the *Pseudomonas* colonies. As shown in Fig. 3A, the two biosensors demonstrated an excellent ability to detect both metabolites, showing orthogonality as expected from Fig. 2F. However, as observed in the top panel of Fig. 3A, the *E. coli* biosensors appear sensitive to the combination of secreted DAPG and pyoluteorin, resulting in growth inhibition of the cells in closest proximity to *P. protegens* DTU9.1.

**Figure 3.**
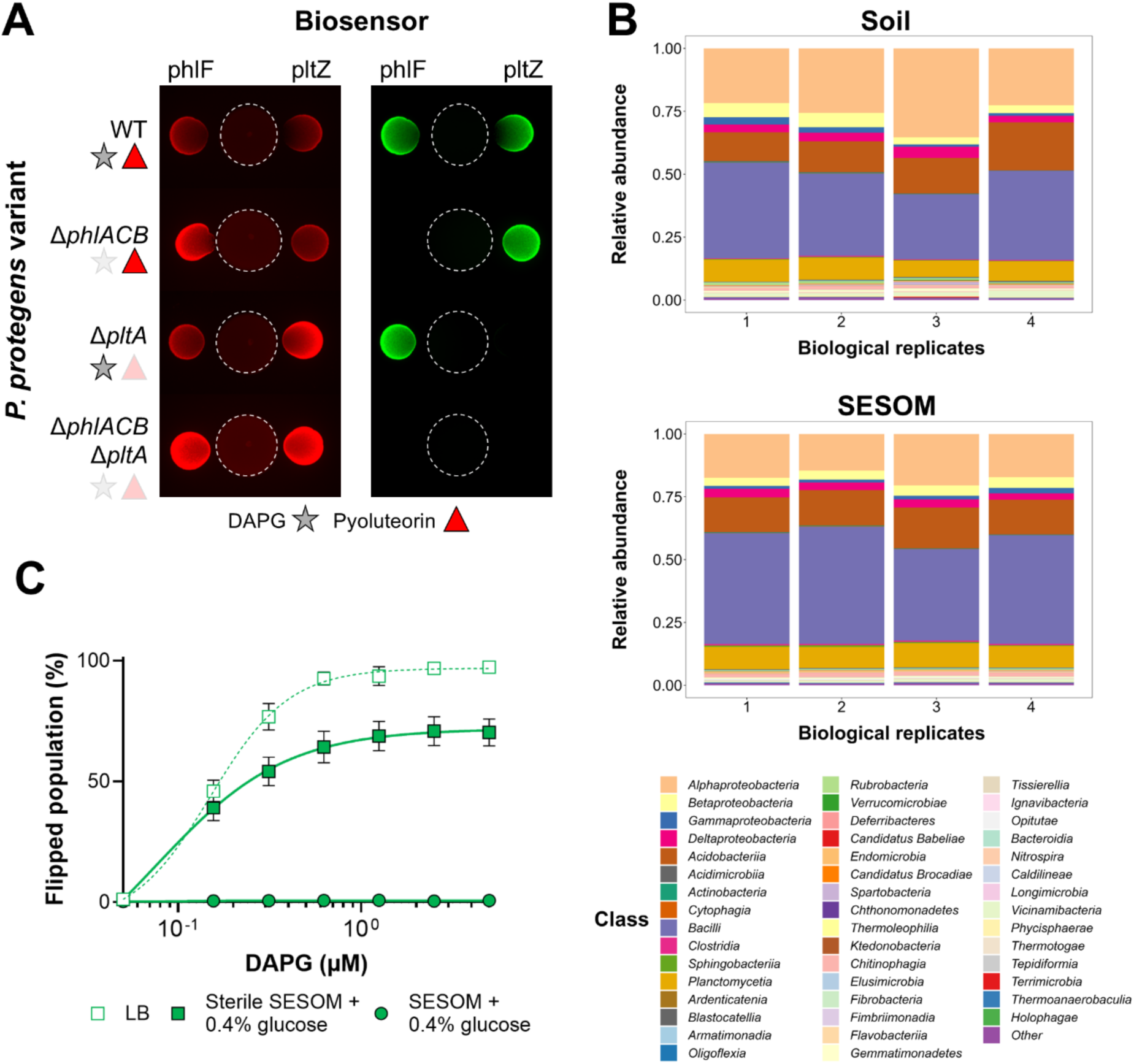
Functionality of *E. coli* biosensors. **A)** Detection of live-produced DAPG and pyoluteorin. *P. protegens* DTU9.1 (indicated by white dotted circle) wildtype and knockout mutants deficient in DAPG and/or pyoluteorin production was pre-cultivated for 24 hours on LB agar before adding *E. coli* biosensors containing either pSEVA231-Int12_LVA_-phlF (*left*) or pSEVA231-Int12_LVA_-pltZ_RBS0.05x_ (*right*). Image stacks were processed with ImageJ and each pair (red and green channel) is a representative of three biological replicates showing similar results. **B)** Comparing bacterial diversity on class level in soil and SESOM. Full-length 16S rRNA gene sequencing was performed using Oxford Nanopore technology on eDNA extracted from four replicates of soil and SESOM prepared from the corresponding soil. **C)** Response curves comparing the DAPG biosensor functionality in sterile and non-sterile SESOM supplemented with glucose compared with LB (see also Supplementary Figure S2). The left-most data point represents the recombination in the absence of inducer.

The catalogue of biosensors was designed to function in soil and rhizosphere environments responding to bioavailable metabolites in complex microbiomes. Thus, we decided to utilize a complex media containing <u>s</u>oil-<u>e</u>xtracted <u>s</u>olubilized <u>o</u>rganic and inorganic <u>m</u>atter (SESOM) as liquid proxy for a soil-like environment (38), which additionally include soil-inhabiting microbes smaller than 10 µm in size. One clear advantage of SESOM for testing our biosensors is its liquid environment, thereby nullifying the possibility of metabolites binding to soil particles, thus becoming unavailable for detection. Initially, we examined the bacterial diversity of SESOM compared to the original soil from which it was extracted to examine if SESOM was a suitable complex environment retaining large parts of the bacterial soil microbiome (Fig. 3B). We extracted eDNA from soil and SESOM and performed full-length sequencing of the 16S rRNA gene using Oxford Nanopore technologies to improve phylogenetic accuracy down to species-level. As shown in Fig. 3B the bacterial diversity of SESOM and the original soil appeared similar, although performing a PERMANOVA indicated that the relative abundance of bacteria classified on the class-level in soil and SESOM was slightly, but significantly different (*P* = 0.032). Nonetheless, SESOM was confirmed to retain a complex bacterial microbiome and would thus serve as a good representation of the soil microbiome, which our biosensors could encounter and interact with during *in situ* detection.

Next, we investigated the functionality of the *E. coli* DAPG biosensor in a sterile and non-sterile version of SESOM (Fig. 3C). The medium was supplemented with 0.4% glucose to mimic the nutrient-scarce rhizosphere environment containing various carbon-rich root exudates (39, 40). As shown in Fig. 3C, the sensitivity of the sensor was retained in sterile SESOM as compared to nutrient-rich LB. However, the sensor was unable to sense and respond to DAPG in non-sterile SESOM. This lack of response was not due to the absence of the *E. coli* biosensor, as it was detectable in both the sterile and non-sterile version of SESOM, although counts of the DAPG biosensor (measured as events of red fluorescing particles) were 10-fold lower in the non-sterile background (Fig. S2). This could indicate that interactions with the microbiome either inhibit parts of the genetic circuit or the *E. coli* host organism. In addition, as mentioned above, specific antimicrobial metabolites (*e.g.* DAPG and pyoluteorin) also prove toxic to *E. coli* at higher concentrations. Thus, *E. coli* might not be the ideal host organism for the genetic memory devices for detection in soil microbiomes.

### Transferring the systems to the soil-inhabiting *Pseudomonas putida*

As an alternative host organism to *E. coli*, *Pseudomonas putida* KT2440 was chosen, since it was originally isolated from soil (41), and we previously identified members of the *P. putida* group to be present in the grassland soil from which we extracted soil and SESOM (14). Thus, we hypothesized that *P. putida* KT2440 would be a suitable host for our genetic memory devices and allow detection of metabolites within complex microbiomes. Martínez-García et al. identified and removed four prophage regions in the genome of *P. putida* KT2440 (42), resulting in the Δall-Φ derivative. We chose this strain as chassis for our genetic circuits to avoid potential, yet uncharacterized interactions between elements of the prophage regions and Integrase12, as this enzyme itself originates from a bacteriophage (29). The two-plasmid system from *E. coli* was altered to chromosomally integrated units (Fig. 4A). The reporter unit was inserted between two convergently transcribed genes PP_1035 and *mltF*, which has previously been characterized as an optimal chromosomal insertion locus (43). The sensory unit containing the metabolite-specific regulator and the *integrase12* gene was inserted upstream of the *glmS* gene as a Tn7 transposon to allow effortless tuning of the various systems.

**Figure 4.**
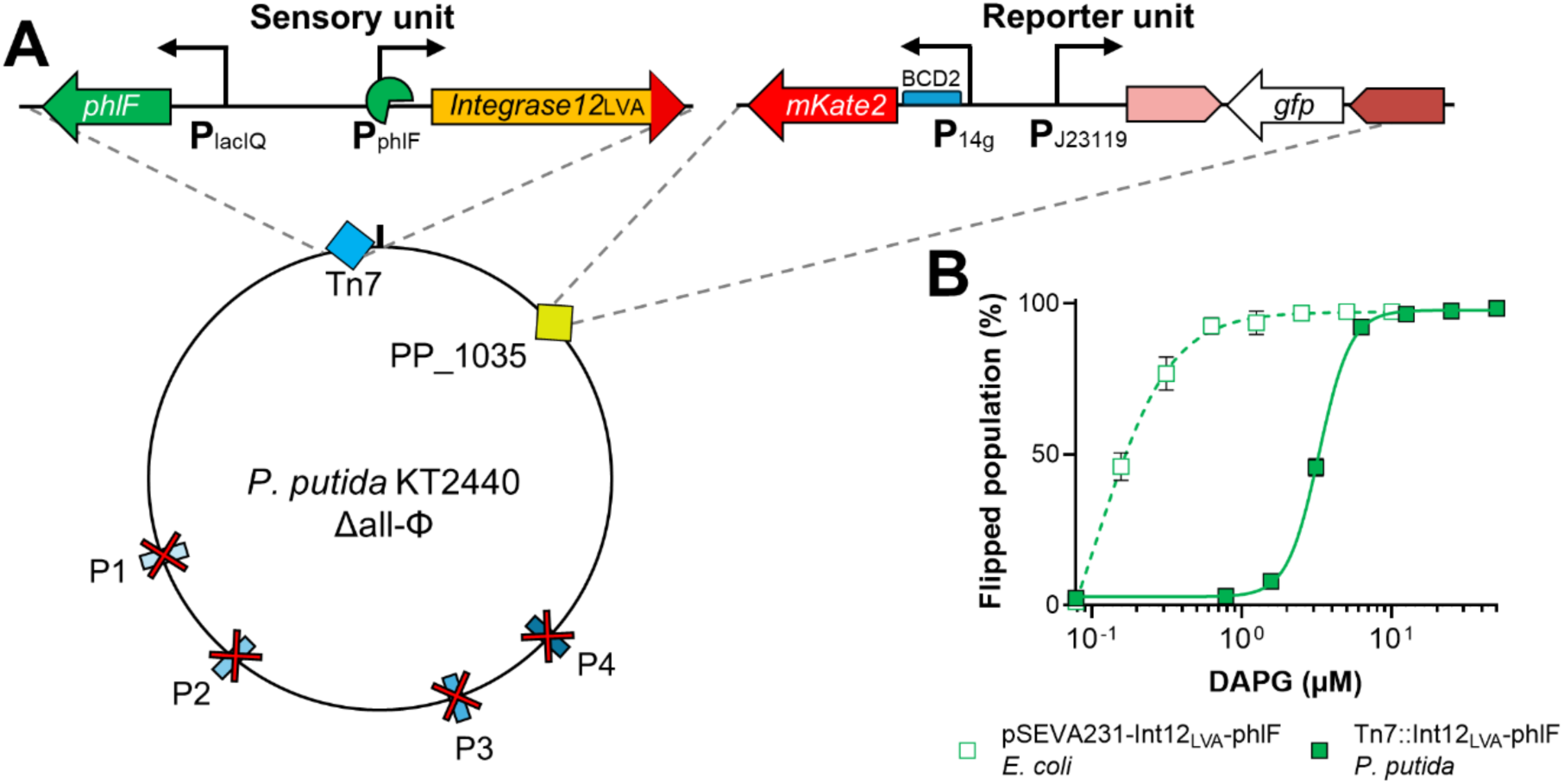
Transferring the genetic memory device to *P. putida*. **A)** The two-unit system was chromosomally integrated in *P. putida* KT2440 Δall-Φ. The sensory unit consisting of the respective regulator and *integrase12* was integrated via Tn7 transposon insertion. Here, the PhlF system is shown as an example. The reporter unit was first changed to constitutively express mKate from the P_14g_ promoter and subsequently integrated downstream of PP_1035 via homologous recombination. **B)** Response curves comparing the sensitivity of the *P. putida* sensor in LB in response to DAPG compared to the original *E. coli* sensor. The left-most data point represents the level of recombination in the absence of inducer.

Testing the DAPG sensing system in *P. putida* KT2440 Δall-Φ showed similar dynamic range as in *E. coli* without undesirable leaky behavior (Fig. 4B), indicating that the protein degradation tag functioned in *P. putida*, which was also shown previously by Andersen et al. (32). However, the sensitivity was 10-fold lower in *P. putida* compared to *E. coli*. This decrease in sensitivity could be attributed to the system being codon optimized and tuned towards functionality in *E. coli*. Nonetheless, the sensitivity remained in the micromolar range, which has previously been shown to reflect concentrations observed in the plant rhizosphere (44).

### Changing the output of the biosensors to genetic memory

In order to utilize the biosensors *in situ* and gain both spatial and temporal resolution of metabolite detection, the output was altered from a fluorescence-based to a DNA-based system, where each biosensor would have a corresponding unique 50 bp barcode located between the *attB*/*P* recognition sites downstream of PP_1035. This design would allow for quantification of cells that had undergone integrase-mediated recombination by qPCR using either Prim1 or Prim2 primers in combination with one of the barcode-specific primers (Fig. 5A).

**Figure 5.**
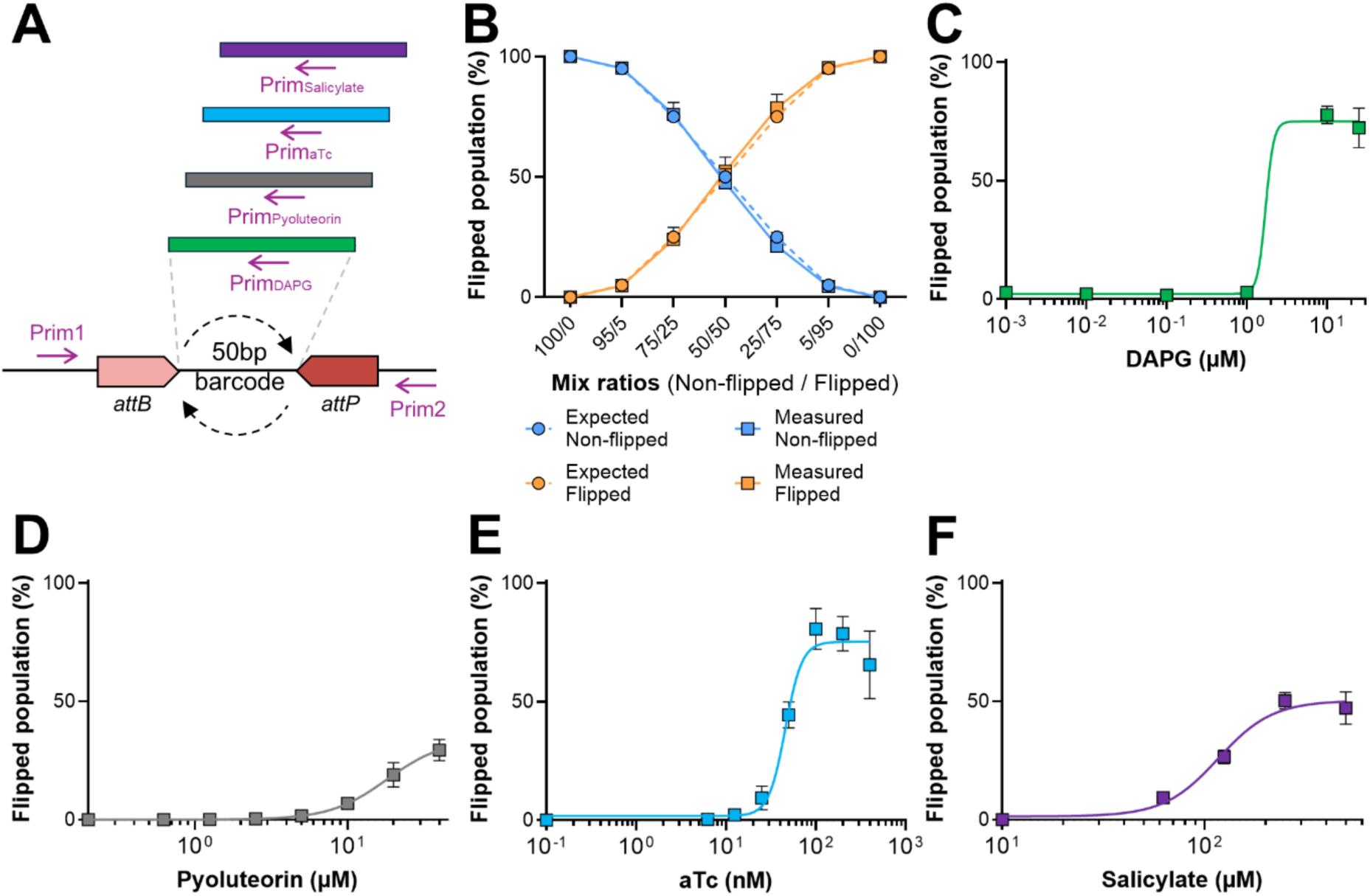
Changing the output to genetic memory. **A)** Schematic representation of the genetic memory output in *P. putida* KT2440 Δall-Φ. The chromosomal reporter unit upstream of PP_1035 was changed to contain only the *a2B/P* recognition sites surrounding a 50bp DNA barcode unique to each sensor. Additionally, primers were designed to amplify short fragments compatible with qPCR, where each DNA barcode had an associated primer which pairs with Prim1 or Prim2 depending on the orientation of the barcode. **B)** Expected versus measured quantities of cells with flipped or non-flipped DNA barcode. *P. putida* KT2440 Δall-Φ strains were constructed to contain the reporter unit with a flipped or non-flipped of the DNA barcode without the sensory unit at the Tn7 site (see Methods section). Cells were mixed in various ratios and cultivated for 24 hours before extracting gDNA and determining absolute quantities of flipped and non-flipped cells with qPCR. **C-F)** Response curves of the genetic memory devices with respective sensory units integrated with Tn7 transposons; DAPG (**C**), pyoluteorin (**D**), aTc (**E**), and salicylate (**F**). Absolute quantities of flipped and non-flipped DNA barcodes in genomic DNA were determined by qPCR using standard curves for each used primer pair. The left-most data points represent the recombination in the absence of inducer.

Initially, we sought to investigate the accuracy of using qPCR to distinguish cells with a native, non-flipped configuration of the DNA barcode from cells with a flipped barcode as a consequence of integrase-mediated recombination. To this end two strains of *P. putida* KT2440 Δall-Φ were cloned with either the non-flipped (attB/P-DAPG barcode_PP_1035_) or flipped (attL/R-**flipped**-DAPG barcode_PP_1035_) barcode specific to the DAPG biosensor, but lacking the sensory unit. These strains were mixed in pre-determined ratios and cultured for 24 hours prior to DNA extraction followed by quantification of non-flipped and flipped cells. As shown in Fig. 5B, the measured quantity of non-flipped and flipped cells accurately matched the predicted ratios, suggesting that genetic memory based on short, unique DNA sequences should provide an accurate and burden-free alternative to the fluorescence-based output. Thus, four biosensors were cloned for detection of DAPG, pyoluteorin, tetracycline, and salicylic acid by chromosomally integrating the respective sensory units with Tn7-based transposition. The naringenin sensor was omitted, as *P. putida* is known to contain the TtgABC eflux pump, whose expression is upregulated by antibiotics and flavonoids (45), suggesting that *P. putida* KT2440 would be a suboptimal host for detection of the flavonoid, naringenin. However, for the remaining four metabolites, the sensors maintained a functional dose-response (Fig. 5C-F), albeit with a reduced dynamic range compared to the fluorescence-based system. This could indicate that integrase-mediated recombination could be affected by the distance between *attB* and *attP*, as recombination requires the formation of a DNA loop aligning the two attachment sites (46). Thus, Integrase12 may be more efficient at recombining the *attB*/*P* pair, when they are distanced by 700-800 nucleotides (presence of *gfp*) compared to the 50-bp DNA barcode. This could suggest that further tuning may be necessary to obtain functional biosensors with DNA-barcode-based output.

### Testing the functionality of the *P. putida* biosensor in soil-like conditions

As the original reason for changing host organism from *E. coli* to *P. putida* was to obtain a chassis that permits metabolite detection in soil-like conditions, an obvious next step was to test the biosensors in SESOM. To this end, the functionality of the fluorescence-based DAPG sensor was evaluated in sterile SESOM supplemented with proline to mimic the availability of root exudates (47). Contrary to the *E. coli* sensor (Fig. 3C), the *P. putida* sensor failed to respond to DAPG in sterile SESOM but performed well LB (Fig. 6A). Thus, we sought to investigate three hypotheses that could explain the lack of functionality of the biosensor. I) The *P. putida* cells were incapable of sensing DAPG, resulting in absence of transcription from the **P**_phlF_ promoter. II) The integrase-mediated recombination was inhibited in *P. putida* under the nutrient-limited conditions in SESOM. III) The stressful conditions of soil-like conditions result in increased ClpXP protease activity, causing an undesirably quick degradation of the tagged Integrase12 before recombination of *attB*/*P* could occur. To test the first hypothesis, *P. putida* KT2440 Δall-Φ containing the pSEVA227-**P**_phlF_-*gfp* was cultivated in sterile SESOM with various concentrations of DAPG. As displayed in Fig. 6B, the cells could indeed sense DAPG, relieve PhlF-repression, and transcribe *gfp* from the **P**_phlF_ promoter.

**Figure 6.**
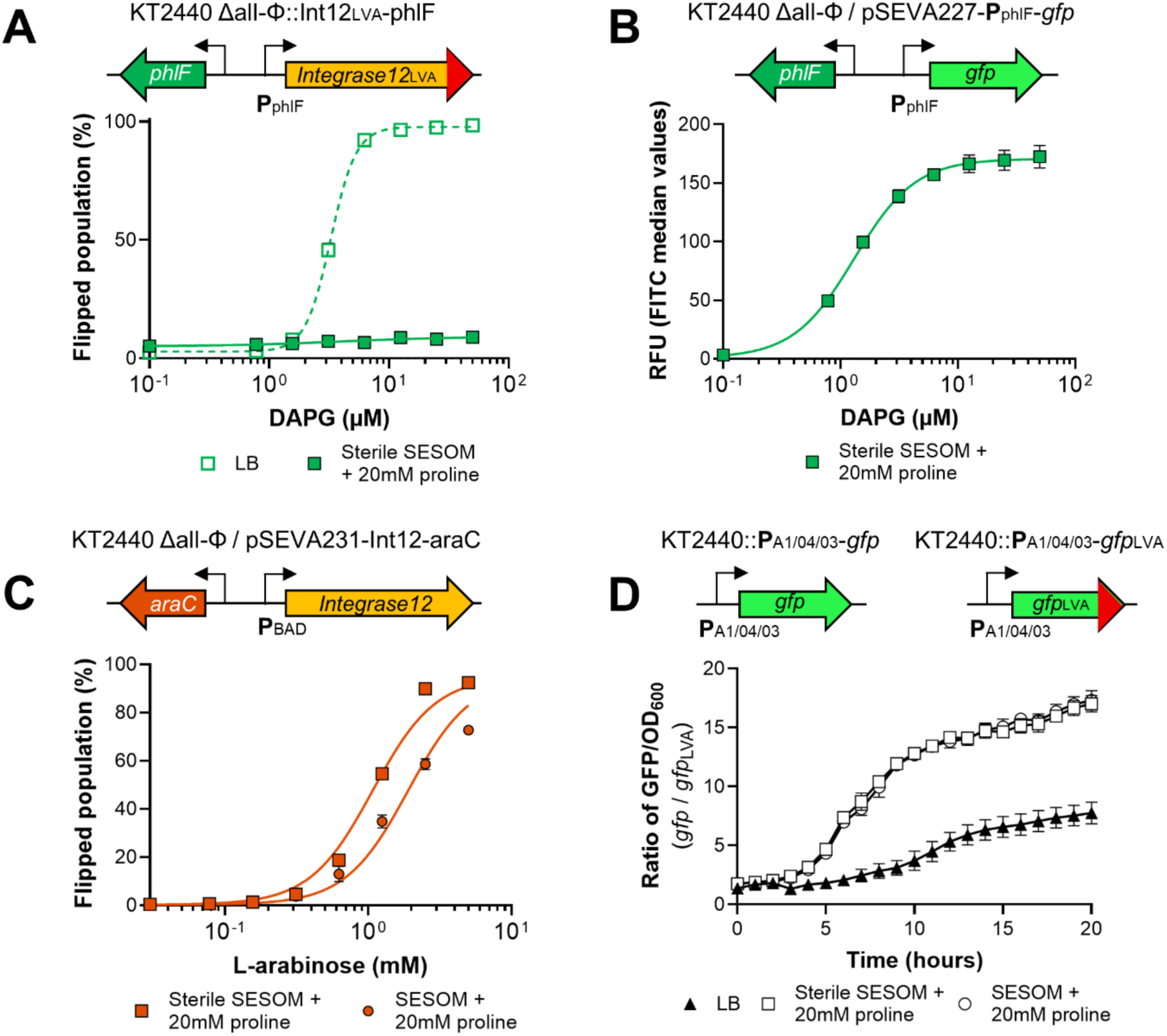
Instability of the integrase makes the system non-functional in a soil-like environment. **A)** Response curves of *P. putida* KT2440 Δall-Φ with the DAPG sensing system in LB and sterile SESOM supplemented with proline and varying concentrations of DAPG. **B)** Response curve of *P. putida* KT2440 Δall-Φ containing pSEVA227-P_phlF_-*gfp* based on relative fluorescence units (RFU) in sterile SESOM supplemented with proline and varying concentrations of DAPG. **C)** Response curves of *P. putida* KT2440 Δall-Φ containing the chromosomal reporter unit and pSEVA231-Int12-araC in sterile and non-sterile SESOM supplemented with proline and varying concentrations of L-arabinose. The left-most data points represent the recombination in the absence of inducer. **D)** Ratio of normalized fluorescence of *P. putida* KT2440 with P_A1/04/03_*gfp* and P_A1/04/03_*gfp*_LVA_ cultivated in LB, sterile SESOM, and SESOM supplemented with proline (see also Supplementary Figure S3).

To test the second hypothesis, we utilized the arabinose-sensor from Yang et al. (29) on the pSEVA231 plasmid. This system was tuned with a difference in copy-number of the sensory and reporter system in mind, as well as the absence of a protein degradation tag. More specifically, in our setup the reporter unit was chromosomally integrated, while the sensory unit on pSEVA231 with the BBR1 origin of replication retained 30-40 copies per *P. putida* cell (48). Thus, this setup could be utilized to investigate if the nutrient-limited environment in SESOM supported the functionality of Integrase12. We expected the system to be non-leaky with a high dynamic range, as shown by Yang et al. (29). Both in sterile and non-sterile SESOM, the arabinose-sensor showed funtionality (Fig. 6C). This finding indicates that integrase-based genetic recombination functions in soil-like conditions, thereby discarding the second hypothesis.

To test the final hypothesis, we utilized the two *P. putida* KT2440 strains from Andersen et al. (32) containing chromosomally integrated *gfp* or *gfp*_LVA_ controlled by the **P**_A1/04/03_ promoter. These strains were cultivated in LB, as well as sterile and non-sterile SESOM supplemented with proline, and their fluorescence was determined over time. To compare the protein degradation rates between different growth media, the ratio of normalized fluorescence of strains containing wildtype *gfp* and *gfp*_LVA_ was determined. As shown in Fig. 6D, the degradation rate of Gfp was increased in both SESOM types compared to LB. Furthermore, the unstable *gfp*_LVA_ variant barely reached fluorescence levels higher than the background in SESOM (Fig. S3). This suggests that the protein degradation rate was increased under the stressful conditions in SESOM to levels causing undesirably rapid degradation of Integrase12, resulting in non-functional biosensors.

### Removal of the degradation tag resulted in functional biosensors in soil-like conditions

In order to further test the hypothesis that the strong degradation tag on *integrase12* resulted in non-functional biosensors under the stressful conditions of SESOM, the LVA tag was removed and the RBS upstream of *integrase12* tuned down to complement the effects of lacking protein instability. One caveat with this approach, and the reason we opted for protein instability in the first place, was that the original Int12-RBS was already tuned low by Yang et al. (29), leaving a very narrow tuning space for decreasing protein translation rate, while retaining predictability. However, using the DAPG biosensor for tuning, an optimal RBS variant was identified, which yielded an almost non-leaky biosensor, although with a smaller dynamic range compared to the sensors relying on Integrase12 instability (Fig. S4).

This RBS was implemented in the four biosensors in *P. putida* KT2440 Δall-Φ and tested for their response to DAPG, pyoluteorin, anhydrotetracycline, and salicylate in sterile and non-sterile SESOM supplemented with proline. As displayed in Fig. 7, all biosensors responded to their respective metabolite under both sterile and non-sterile soil-like conditions supplemented with proline. The DAPG biosensor appeared leaky in the absence of DAPG (Fig. 7A). However, none of the biosensors demonstrated undesirable integrase-mediated recombination prior to starting the assays in SESOM (Fig. S5), suggesting that the observed leakiness could in fact be a response to DAPG present in the SESOM. Although, we were unable to detect DAPG in SESOM using LCMS/MS on extracted metabolites utilizing an approach previously used to detect DAPG in a liquid environment (8), we have in fact previously isolated and identified DAPG-producing bacteria from the same soil used to prepare SESOM (14). Importantly, the full-length 16S rRNA profiling of the SESOM revealed the presence of *Pseudomonas* species belonging to the *P. fluorescens* group, which is known to contain DAPG producers (49), suggesting that the biosensor responded to DAPG, or other similar metabolites already present in SESOM capable of relieving PhlF-mediated repression. Overall, the results from the SESOM experiments indicate that the engineered catalogue of biosensors in *P. putida* KT2440 Δall-Φ should be functional in a rhizosphere environment for *in situ* detection within complex microbiomes.

**Figure 7.**
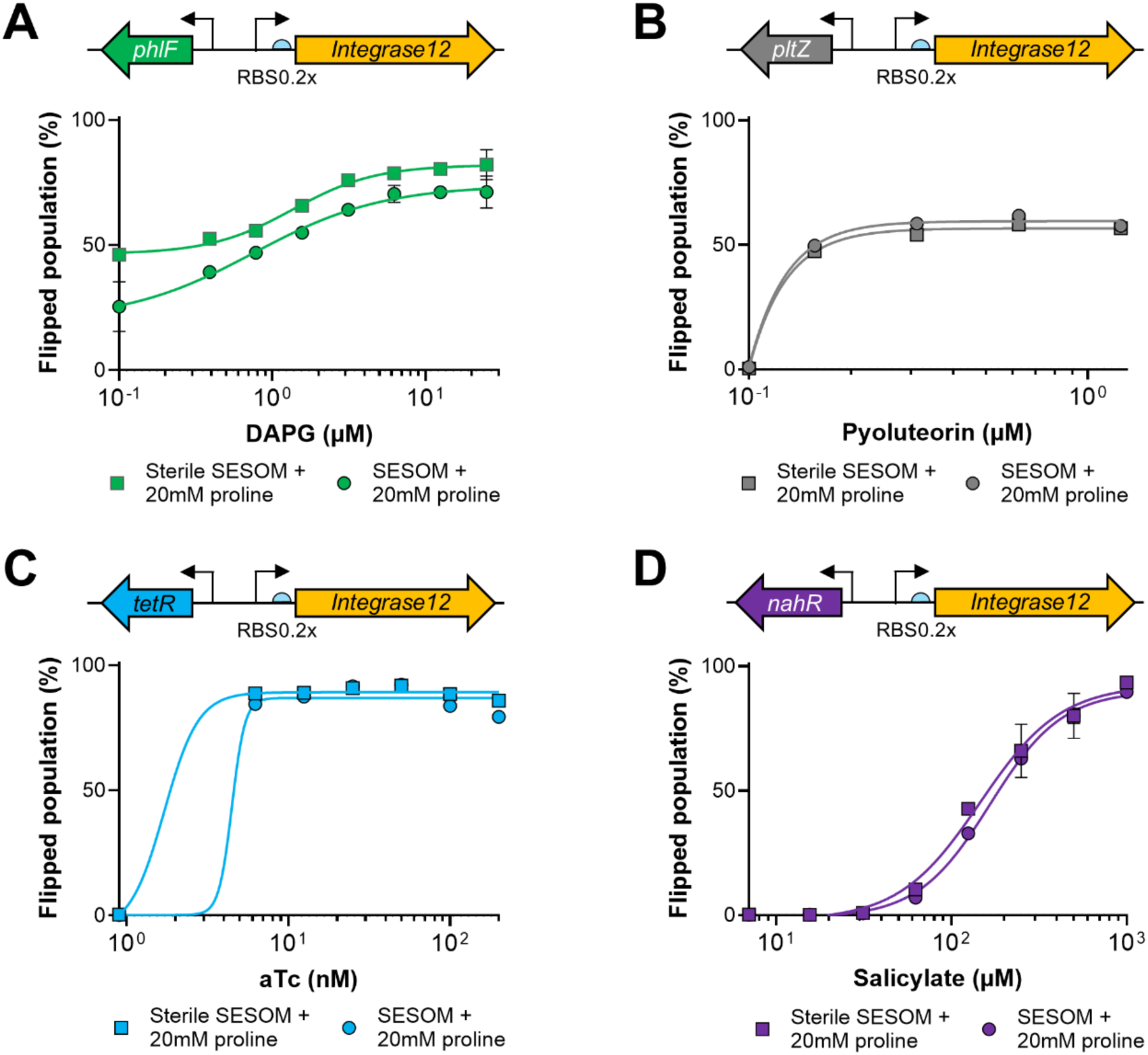
Genetic memory devices functioning in soil-like conditions. Response curves for the optimized biosensors without degradation tag on *integrase12* and Int12-RBS0.2x in response to DAPG (**A**), pyoluteorin (**B**), anhydrotetracycline (**C**), and salicylate (**D**) in SESOM and sterile SESOM supplemented with proline (see also Supplementary Figure S5). The left-most data points represent the recombination in the absence of inducer.

### Perspectives of genetic memory devices

Genetic memory devices as presented here have a clear advantage over the common transcription factor-based biosensors with a fluorescent or colorimetric output in that these “sense and record”-devices with DNA barcode-based output can be mixed in cocktails permitting multiplexed detection of various analytes simultaneously. The uniqueness of the DNA barcodes allows for parallelized experiments, as readouts can be easily distinguished with amplicon sequencing of extracted environmental DNA covering the *attB*/*P* region. An additional benefit of DNA-based output is the absence of the cellular burden posed by the fluorescence-based reporter system, which should permit stability of the recording devices in long duration *in vivo* experiments. Kalvapalle et al. demonstrated that considering the cellular burden is critical to obtain stable biosensor populations over time, as they opted for a frugal setup, where induced integrase-mediated recombination excised the entire biosensor system, thereby nullifying the burden of any output system, as well as *integrase* expression in the presence of analyte after recombination had taken place (50). Such a system may be the ideal burden-free design, although, it is highly prone to leaky expression, resulting in heterologous cell populations with non-functional biosensors. Our design utilizing unique DNA barcodes as readout (Fig. 5) may therefore represent a suitable alternative, which is easier to clone and maintain, as well as scale up to include sensing of multiple different analytes simultaneously.

Single-component transcription factors are the most abundant natural sensors (51) and have evolved over billions of years to regulate cellular responses to environmental changes and stimuli. Thus, a large repertoire of known transcription factors and their cognate ligand are available (51–53), which could effortlessly be implemented in the memory device chassis proposed here, thereby further broadening the catalogue of metabolites, signals, and other mediators of microbial communication that can spatiotemporally be sensed *in situ*. Additionally, bioinformatic tools enable the prediction of novel regulators and their associated operator based on ligand structure, thereby providing the opportunity of constructing novel biosensors (54, 55).

Monitoring transient metabolites and other microbial signals *in situ* without disrupting experimental setups and existing microbiomes is an admiral goal of whole-cell biosensors and will provide much needed information about the chemical dialogue occurring constantly between plants and microbes. However, the concept of “sense and record” may also prove as a steppingstone for more sophisticated “sense and deliver” systems. For instance, biosynthesis of the plant hormone salicylic acid is induced by pathogen invasion, which in turn induces systemic resistance as a broad-spectrum response to combat infection (24). As a result, salicylic acid concentration increases in root exudates (25, 26). Thus, our root-colonizing salicylate biosensor could be further expanded upon by coupling sensing to biosynthesis of a toxin specific to the particular plant pathogen and thereby aid in effective and local biocontrol as response to the plant’s salicylate-associated “cry for help”. Others have already demonstrated the potential application of such engineered, biological devices as tools to fixate nitrogen (56) and promote plant growth (57).

## Conclusion

Here we constructed a catalogue of five whole-cell biosensors for spatiotemporal, non-disruptive *in situ* detection of biologically relevant specialized plant and microbial metabolites. Four of these biosensors were successfully transferred to the soil-compatible *P. putida* KT2440 Δall-Φ strain. Lastly, we demonstrated the functionality of the sensors in a soil-extracted medium containing the diverse microbiome found in soil. One of the key advantages of genetic memory devices is the ability to monitor bioavailable, transient signals utilized in the chemical dialogue between plants and microbes. Additionally, the chassis of biosensors with DNA-barcodes as readout proposed here is readily scaled to sense more analytes in the future.

## Methods

### Strains and media

*E. coli* TOP10 and CC118λ*pir* were used for genetic manipulation, while *E. coli* DH10B and *P. putida* KT2440 Δall-Φ were used for genetic circuit characterization. Cells were routinely grown in LB Miller broth (CarlRoth, X964) for cloning, as well as functional assays unless otherwise stated. For agar plates 2% w/v food-grade agar was supplemented (PanReac AppliChem, A0917). For sucrose selection, NSLB agar plates were used (10 g/L tryptone, 5 g/L yeast extract, 15 g/L Bacto agar) supplemented with 15% v/v sucrose (Sigma-Aldrich, 84100). Chloramphenicol (Sigma-Aldrich, C0378), kanamycin (Sigma-Aldrich, 60615) or gentamicin (Sigma-Aldrich, G1264) was supplemented where appropriate. 2,4-diacetylphloroglucinol (Apollo Scientific, OR22030), pyoluteorin (Santa Cruz Biotechnology, SC-391693A), anhydrotetracycline (Sigma-Aldrich, 37919), salicylic acid (Sigma-Aldrich, 105910), naringenin (Sigma-Aldrich, N5893), and L-arabinose (Sigma-Aldrich, A3256) were used as inducers.

### Preparation of Soil-Extracted Solubilized Organic and inorganic Matter (SESOM)

Grassland soil was collected from sampling site P8 in Dyrehaven, Denmark (55⁰79’52”N, 12⁰58’06”E). The soil was initially sifted through a 2 mm sifting tray. Then, 80 g of sifted soil was added to 400 mL of 1X PBS and incubated at 30°C shaking at 180 rpm for 3 hours. The soil slurry was run through a triple-layered miracloth (VWR, 475855-1) to remove larger soil particles. Then, the flowthrough was filtered first through double-layered 25 µm Whatman filter (Sigma-Aldrich, WHA1004125) in a Büchner funnel/vacuum setup and secondly through a double-layered 11 µm Whatman filter (Sigma-Aldrich, WHA1001090). Finally, the flowthrough was filtered using a 10 µm Omnipore PTFE membrane (Sigma-Aldrich, JCWP04700) using a Rocker 300 – SF30 vacuum filtration system to generate SESOM. Sterile SESOM was produced by running a subsample of SESOM through 0.22 µm filter.

### Cloning of sensor-integrase plasmids

Integrase plasmids (pInt12 and pSEVA231 variants, see Supplementary Table 3) were cloned using Golden Gate cloning with high-fidelity enzymes from New England Biolabs. Promoters, RBS sequences, and protein degradation tags were introduced with primers synthesized by IDT DNA Technologies. Libraries of varying RBS strengths measured as translation initiation rates relative to the respective host organism were predicted with RBScalculator (36). Regulators were either cloned from pAJM.847 (*phlF*), pAJM.011 (*tetR*), pAJM.771 (*nahR*), pInt12-araC (*araC*) or synthesized as *E. coli* codon optimized gBlocks from IDT DNA Technologies (*pltZ* and *fdeR*). All genetic parts can be found in Supplementary Table 5. Cloned plasmids were transformed into chemically competent *E. coli* TOP10 cells and transformants selected on LB supplemented with 50 µg/mL kanamycin. Cloned plasmids were verified by Sanger sequencing at Eurofins Genomics. *E. coli* DH10B cells containing pRep12 were made chemically competent and transformed with integrase plasmids containing the different regulators. Transformants were selected on LB agar supplemented with 50 μg/mL kanamycin and 10 μg/mL chloramphenicol.

Plasmids designed for Tn7 transposon insertion in *P. putida* KT2440 Δall-Φ (see Supplementary Table 3) were cloned with classic restriction cloning using SacI (ThermoFisher, FD1133) and XbaI (ThermoFisher, FD0684), where the entire regulator/integrase part from pSEVA231 variants was cloned into the pTn7-M vector. Cloned pTn7 plasmids were transformed into chemically competent *E. coli* CC118λ*pir* cells and selected on LB supplemented with 10 µg/mL gentamicin. Cloned plasmids were verified by Sanger sequencing at Eurofins Genomics.

### Triparental and four-parental mating

Constructs in pSEVA backgrounds replicable in *Pseudomonas* were conjugated into *P. putida* KT2440 Δall-Φ through tri-parental mating with *E. coli* CC118λ*pir* and *E. coli* HB101 / pRK600, which encodes the RP4 conjugative machinery. Constructs in the pTn7-M background were conjugated into *P. putida* KT2440 Δall-Φ through four-parental mating with *E. coli* CC118λ*pir*, *E. coli* HB101 / pRK600, and *E. coli* Pir1 / pTNS2, which encodes the transposase genes. Conjugation mixes were prepared by washing O/N cultures of implicated strains in 1X PBS once, followed by adjusting OD_600_ to 1.0, mixing cultures in equal volumes in a total of 1 mL. Culture mixes were centrifuged at 10,000 rpm for 1 minute, followed by discarding the supernatant by decanting, resuspending the pellet in the remaining liquid, and incubating it as a spot on LB agar at 30°C O/N. Transconjugants were selected on *Pseudomonas* isolation agar (Sigma-Aldrich, 17208) supplemented with appropriate antibiotics.

### Cloning and integrating reporter units in *P. putida* KT2440 Δall-Φ

For assays in *P. putida* KT2440 Δall-Φ reporter units containing either a fluorescence-based or DNA-based detection system (Supplementary Table 2) were designed for chromosomal integration upstream of PP_1035 using an allelic replacement approach. The pK18msB plasmid containing a kanamycin resistance cassette for initial selection and a *sacB* gene for secondary sucrose counter-selection was chosen as suicide vector (58). Reporter units were introduced between two convergently transcribed genes PP_1035 and *mltF* (between nt 1,182,901 and 1,182,902). For DNA-based detection, five unique 50 bp barcodes were chosen from pMemoryArray (29). DNA barcodes and the two 500 bp homology regions up- and downstream of the insertion site at PP_1035 (HR1_PP_1035_ and HR2_PP_1035_, see Supplementary Table 5) were cloned into pK18msB using Golden Gate assembly. Additionally, a flipped version of the DNA-based system with the DAPG-barcode was cloned by designing the attL/R-**flipped**-DAPG barcode *in silico* and introducing it into pK18msB along with the homology regions using primers with extended overhangs. For fluorescence-based detection, the origin of replication of pK18msB plasmid was initially changed to R6K to obtain a low-copy version allowing for cloning of burdening system in *E. coli*, such as the continuously high-expression **P**_14g_-*mKate* system. To obtain pK18msB_R6K_-Rep12_PP_1035_ the attB/P-*gfp* system from pRep12, the **P**_14g_-*mKate* from pBG42-mKate, as well as the two homology regions (HR1_PP_1035_ and HR2_PP_1035_) were cloned into pK18msB_R6K_ using Golden Gate assembly. Plasmids were transformed into *E. coli* CC118λ*pir* and transformants selected on LB supplemented with 50 µg/mL kanamycin. Positive clones were verified with Sanger sequencing at Eurofins Genomics.

For chromosomal integration of reporter units in *P. putida* KT2440 Δall-Φ the pK18msB suicide vectors were initially transferred with triparental mating and selected on *Pseudomonas* isolation agar supplemented with 50 µg/mL kanamycin. For counter-selection, a single colony was inoculated into LB O/N at 30°C and dilutions plated on NSLB agar plates. Correct insertion of reporter units was verified with PCR. Tn7 transposons with the sensory units (mentioned above) were subsequently introduced into these strains.

### Characterization of biosensors

To characterize the function of the *E. coli* DH10B biosensors (**Fig. 1B-C**, **Fig. 2** and **Fig. S1**), overnight cultures were prepared in three biological replicates in LB supplemented with 25 μg/mL kanamycin and 10 μg/mL chloramphenicol. Overnight cultures were diluted 1:100 in 200 µL LB containing 25 μg/mL kanamycin and 10 μg/mL chloramphenicol in the presence of various inducers in flat-bottom 96-well plates (VWR, 10062-900) covered with air-permeable Breathe-Easy sealing membranes (Sigma-Aldrich, Z380059) and incubated for 20 hours at 37°C at 600 rpm. To characterize the induction of Integrase12 over time (**Fig. 1D**), cells containing pRep12 and pSEVA231-Int12_LVA_-phlF were diluted 1:100 in 1 mL LB containing 25 μg/mL kanamycin and 10 μg/mL chloramphenicol and induced with 25 µM DAPG in U-bottom 96-deep-well plates (Eppendorf, 951032603) covered with a silicone sealing mat (Corning, AM-2ML-SQ) for 8 h at 37°C at 600 rpm, where 50 µL samples were taken every hour.

To characterize the function of the *E. coli* DH10B DAPG biosensor under sterile and non-sterile soil-like conditions (**Fig. 3C** and **Fig. S2**), overnight cultures of *E. coli* DH10B containing pRep12 and pSEVA231-Int12_LVA_-phlF were prepared in three biological replicates in LB supplemented with 25 μg/mL kanamycin and 10 μg/mL chloramphenicol. Overnight cultures were washed twice in 1X PBS and diluted 1:100 in 200 µL SESOM and sterile SESOM supplemented with 0.4% glucose and varying DAPG concentrations in flat-bottom 96-well plates covered with air-permeable Breathe-Easy sealing membranes and incubated for 20 hours at 37°C at 600 rpm. For flow cytometry analysis, a 25 μL aliquot of each culture in LB was added to 175 μl 0.9% NaCl and a 50 µL aliquot of each culture in SESOM was added to 150 μl 0.9% NaCl.

To characterize the function of the *P. putida* KT2440 Δall-Φ biosensors (**Fig. 4B**, **Fig. 6A** and **6C**, **Fig. 7**, and **Fig. S4**), overnight cultures of *P. putida* KT2440 Δall-Φ Rep12_PP_1035_ with Tn7-inserted sensor units were prepared in three biological replicates in LB supplemented with 25 μg/mL gentamicin. Prior to assaying the biosensors in SESOM and sterile SESOM supplemented with 20 mM L-proline (Sigma-Aldrich, 1.07434), a 20 µL aliquot was taken from overnight cultures, added to 980 µL 0.9% NaCl, and analyzed by flow cytometry to determine the percentage of pre-flipped sensor cells in the absence of inducer (**Fig. S5**). For the assays, overnight cultures were diluted 1:100 in 200 µL LB or SESOM and sterile SESOM supplemented with 20 mM L-proline in the presence of various inducers in flat-bottom 96-well plates covered with air-permeable Breathe-Easy sealing membranes and incubated for 20 hours at 30°C at 600 rpm. For flow cytometry analysis, a 2 μl aliquot of each culture was added to 198 μl 0.9% NaCl.

### Flow cytometry analysis

Fluorescence was measured using a MACSQuant VYB flow cytometer (Miltenyi Biotec). For each sample up to 10^5^ events were recorded using a medium flow rate. FlowJo v10 was used to analyze the data. All events were initially gated by forward scatter and side scatter. Red fluorescence (>10^1^ a.u.) was subsequently used for gating cells containing pRep12 or the chromosomal reporter unit. Events corresponding to negative GFP fluorescence were excluded. The flipped population was based on the percentage of cells above a GFP threshold higher than cells in off-state in the absence of inducer. Graphs were produced with GraphPad Prism 10, and for response curves data was fitted with a dose/response curve to guide the eye.

### Analysis of P_phlF_ activity in SESOM

To assay the activity of the **P**_phlF_ promoter in SESOM (**Fig. 6B**), the pSEVA227-**P**_phlF_*-gfp* plasmid (14) was transferred to *P. putida* KT2440 Δall-Φ by triparental mating and selected on *Pseudomonas* isolation agar supplemented with 50 µg/mL kanamycin. Overnight cultures prepared in three biological replicates were washed twice in 1X PBS and had their OD_600_ normalized. Cultures were inoculated in 200 µL SESOM and sterile SESOM supplemented with 20 mM L-proline and varying concentrations of DAPG to an initial OD_600_ = 0.01. The microplate was sealed with a Breathe-Easy membrane and incubated for 20 hours at 30°C and 600 rpm shaking. Aliquots of 50 µL were added to 150 µL 0.9% NaCl and analyzed by flow cytometry to determine fluorescence intensity.

### Protein-degradation assay

To compare the Clp-protease-dependent degradation rate in LB and SESOM, *P. putida* KT2440::**P**_A1/04/03_*gfp* and KT2440::**P**_A1/04/03_*gfp*_LVA_ were cultured in three biological replicates overnight prior to starting a growth assay in a 96-well microplate (**Fig. 6D** and **Fig. S3**). Overnight cultures were washed twice in 1X PBS and normalized to OD_600_ = 1.0. Cultures were inoculated in 200 µL LB, as well as SESOM and sterile SESOM supplemented with 20 mM L-proline to an initial OD_600_ = 0.01. The microplate was sealed with a Breathe-Easy membrane and incubated for 20 hours at 30°C with continuous shake in a BioTek Synergy H1 multimode reader (Agilent). Readings of OD_600_ and GFP fluorescence were taken every 15 minutes.

### Detection of live-produced DAPG and pyoluteorin

To investigate the functionality of the *E. coli* DH10B biosensors in detection of live-produced metabolites (**Fig. 3A**), *P. protegens* DTU9.1 wildtype, Δ*phlACB*, Δ*pltA*, and the double knockout (Supplementary Table 1) were precultured in three biological replicates in LB prior to normalizing their OD_600_ to 1. Subsequently, 20 µL normalized culture was spotted on LB agar plates, where each variant of *P. protegens* DTU9.1 was spotted on separate plates to avoid unintentional diffusion of metabolites from adjacent colonies. Plates were incubated at 30°C for 24 hours before adding the biosensors. Cultures of *E. coli* DH10B containing pRep12 and either pSEVA231-Int12_LVA_-phlF or pSEVA231-Int12_LVA_-pltZ_RBS0.05x_ were normalized to OD_600_ = 1 and spotted as 5 µL colonies next to the *P. protegens* DTU9.1 colony. Plates were re-incubated at 37°C for 24 hours before microscopic imaging on a Zeiss Axio Zoom.V16 fluorescence microscope (Zeiss). Image stacks were analyzed with ImageJ.

### Characterization of bacterial diversity in soil and SESOM

To compare bacterial diversity in soil and SESOM (**Fig. 3B**), eDNA was extracted with DNeasy PowerSoil Pro kit (Qiagen, 47016) using the manufacturer’s instructions. For soil, eDNA was extracted from 500 mg of soil sifted with a 2 mm sifting tray. For SESOM, eDNA was extracted from 2 mL. Additionally, 50 µL 0.5 M Fe_3_Cl (Sigma-Aldrich, F2877) was added to bead beating tubes along with Lysis Buffer 1 (see DNeasy PowerSoil Pro kit manual) to reduce amounts of PCR inhibitors (59). The 16S rRNA gene was amplified with barcoded primers from the 16S Barcoding Kit 24 V14 (Oxford Nanopore, SQK-16S114.24). For soil, 1 µL 100x diluted extracted eDNA was used as template, while 10 µL undiluted extracted eDNA was used as template for SESOM. PCR and preparation of the sequencing mix was performed using manufacturer’s instructions. The pooled, barcoded 16S rRNA amplicons were sequenced on a Nanopore flow cell (Oxford Nanopore, FLO-MIN114). Reads were base-called with super-high accuracy and taxonomical classification. Reads were trimmed with Chopper (60) and de-barcoded with Porechop (61). Relative species abundance based on full-length 16S rRNA genes was performed with Emu (62). The diversity of soil and SESOM was compared with a PERMANOVA using the Adonis tool in the vegan R package (63).

### Quantitative PCR

To determine the function of the genetic memory devices in *P. putida* KT2440 Δall-Φ using DNA memory as output, qPCR was utilized to quantify the levels of integrase-mediated recombination with primer pairs specific to the non-flipped DNA barcode (Prim1/Prim_barcode_) or flipped DNA barcode (Prim2/Prim_barcode_). Primers are summarized in Supplementary Table 4. To determine the sensitivity of utilizing the qPCR-based quantification method (**Fig. 5B**), three biological replicates of cultures of *P. putida* KT2440 Δall-Φ containing either attB/P-DAPG barcode_PP_1035_ or attL/R-**flipped**-DAPG barcode_PP_1035_ were OD_600_-normalized and mixed in pre-defined ratios. The culture mixes were incubated for 24 hours in LB at 30°C and 200 rpm shaking prior to extracting gDNA by boiling and determining the measured ratios of cells containing non-flipped and flipped DNA barcodes. To investigate the functionality of biosensors using DNA-memory as output (**Fig. 5C-F**), overnight cultures of *P. putida* KT2440 Δall-Φ attB/P-barcode_PP_1035_ with corresponding Tn7-inserted sensor units (Supplementary Table 2) were prepared in three biological replicates in LB supplemented with 25 μg/mL gentamicin. For the assays, overnight cultures were diluted 1:100 in 200 µL LB in the presence of various inducers in flat-bottom 96-well plates covered with air-permeable Breathe-Easy sealing membranes and incubated for 20 hours at 30°C at 600 rpm prior to extracting gDNA by boiling.

For qPCR, template DNA was prepared by adding 15 µL cell culture to 45 µL nuclease-free water and boiling for 20 minutes to extract gDNA, where 2 µL cell lysate was used as template DNA. The qPCR reactions were run with 2X Luna Universal qPCR Master Mix (New England Biolabs, M3003) in 0.2 mL Non-skirted 96-well PCR plates (Thermo Scientific, AB-0600) covered with Optically Clear flat strips (Thermo Scientific, AB-0866) with 0.4 µM of each primer in a final volume of 20 µL. Reactions were run on a Bio-Rad CFX Opus 96 qPCR machine (Bio-Rad), with the following conditions: 95 °C for 2 min and 40 cycles of 95 °C for 15 sec, 60 °C for 30 sec. To obtain a melting curve profile of the qPCR products 60 cycles of 5 sec at 65 °C to 95 C with a 0.5 °C temperature increase during each step was performed. Data was analyzed with CFX Maestro software 2.3 (Bio-Rad). For absolute quantification, we used a standard curve comparing CFU/mL to Cq-value generated for each strain tested (ranging from 10^4^-10^9^ CFU/mL). Flipping ratio was calculated as (CFU/mL of cells with flipped DNA) / (Total CFU/mL of cells with flipped and non-flipped DNA).

## Data availability

Demultiplexed 16S rRNA sequencing reads utilized to profile the bacterial diversity of soil and SESOM were uploaded to NCBI SRA database under BioProject number PRJNA1216427. The tables containing data of relative abundance at species level in both soil and SESOM, as well as maps of cloned plasmids can be found at https://doi.org/10.11583/DTU.28270904.

## Acknowledgement

We thank the members of the Center for Microbial Secondary Metabolites (CeMiSt) for general scientific discussions and advice. We thank Esteban Martínez-García for providing the strain, *P. putida* KT2440 Δall-Φ, as well as Pablo Nikel for providing several pSEVA vectors. The pIntegrase_12 and pReporter_12 plasmids were a gift from Christopher Voigt (Addgene plasmid #60583 and #60572, respectively). This study was carried out with funding from the Villum Foundation (Grant number 40964). This study was additionally funded by The Danish National Research Foundation (DNRF137) as part of CeMiSt. Additionally, funding was received from the Novo Nordisk Foundation (NNF19OC0055625) for the infrastructure “Imaging microbial language in biocontrol (IMLiB)”.

## Notes

**Conflict of Interest:** Authors declare no conflict of interest.

### Competing Interest Statement

The authors have declared no competing interest.

