## Supplementary file for "Genetic memory devices to detect specialized metabolites in plant and soil microbiomes"

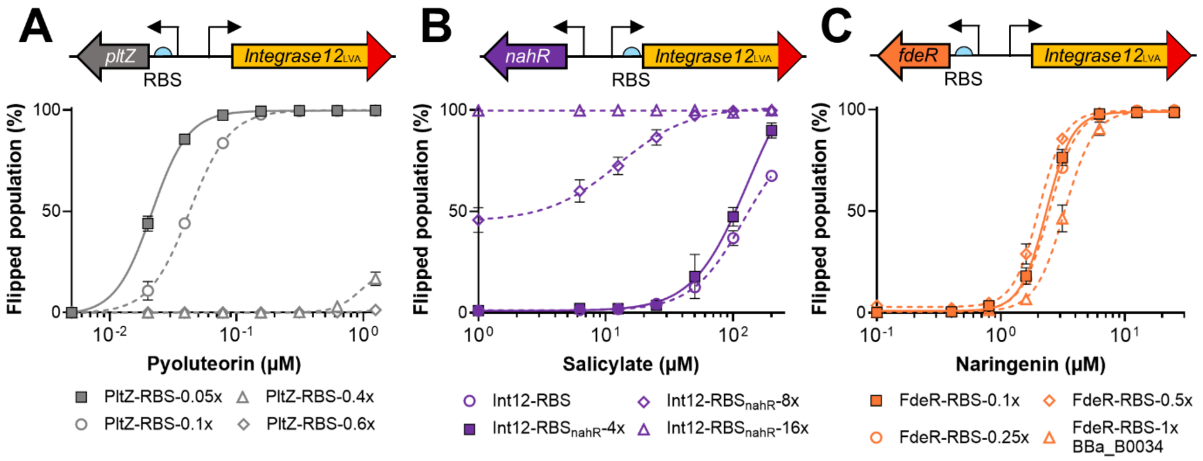


**Figure S1 | Tuning the *E. coli* biosensors.** The biosensors were tested in *E. coli* DH10B containing both pRep12 and pSEVA231-Int12$_{LVA}$ with respective regulators. Sensitivity was tuned for the pyoluteorin, salicylate, and naringenin sensor. For the pyoluteorin (**A**) and naringenin (**C**) sensors, the RBS upstream of the regulator gene was tuned, whereas the RBS upstream of *integrase12* was tuned for the salicylate sensor (**B**). The optimal variants are shown with solid symbols and lines. All RBS sequences are provided in Supplementary Table 5. The left-most data points represent the recombination in the absence of inducer. Data represents the mean of the three biological replicates.

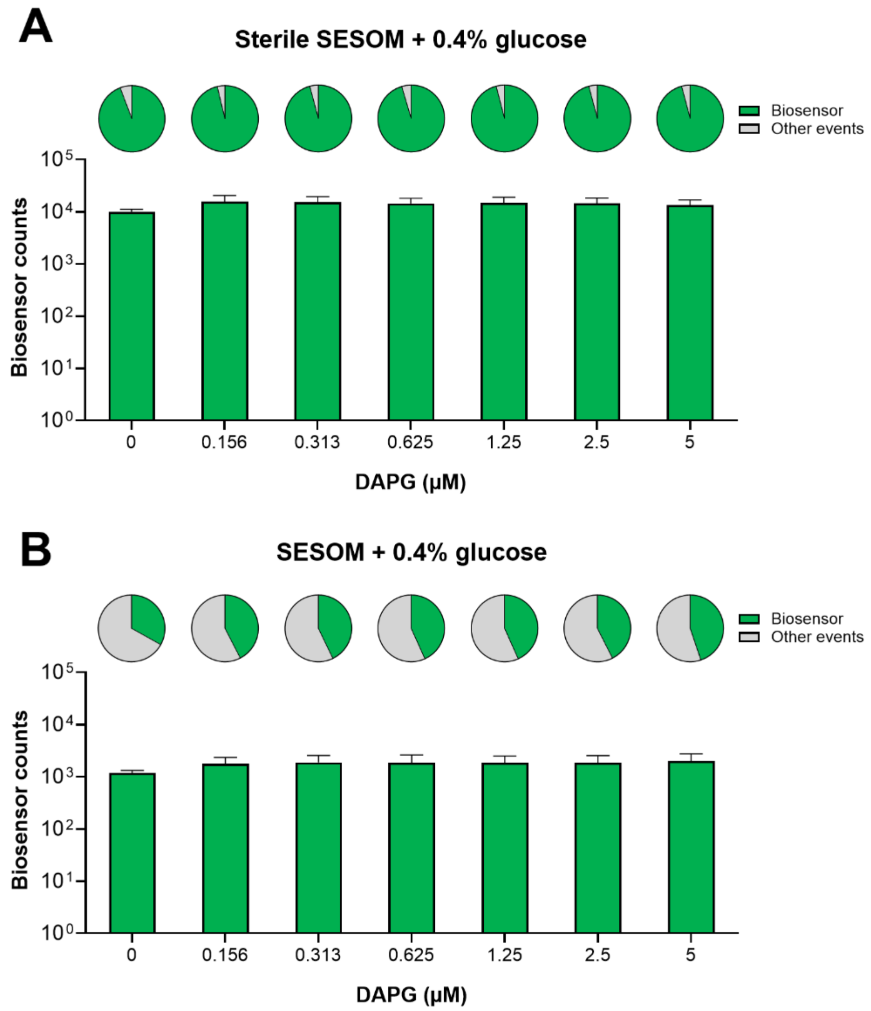


**Figure S2 | Counts of *E. coli* DAPG biosensor in sterile and non-sterile SESOM.** Bars represent counts of red fluorescent cells (biosensors with pRep12) determined by flow cytometry. Data is shown as mean values of three biological replicates with error bars displaying standard deviations. Pie diagrams on top represent the average percentage of biosensor cells compared to the total amount of events for each concentration of DAPG tested in sterile (**A**) and non-sterile (**B**) SESOM supplemented with 0.4% glucose.

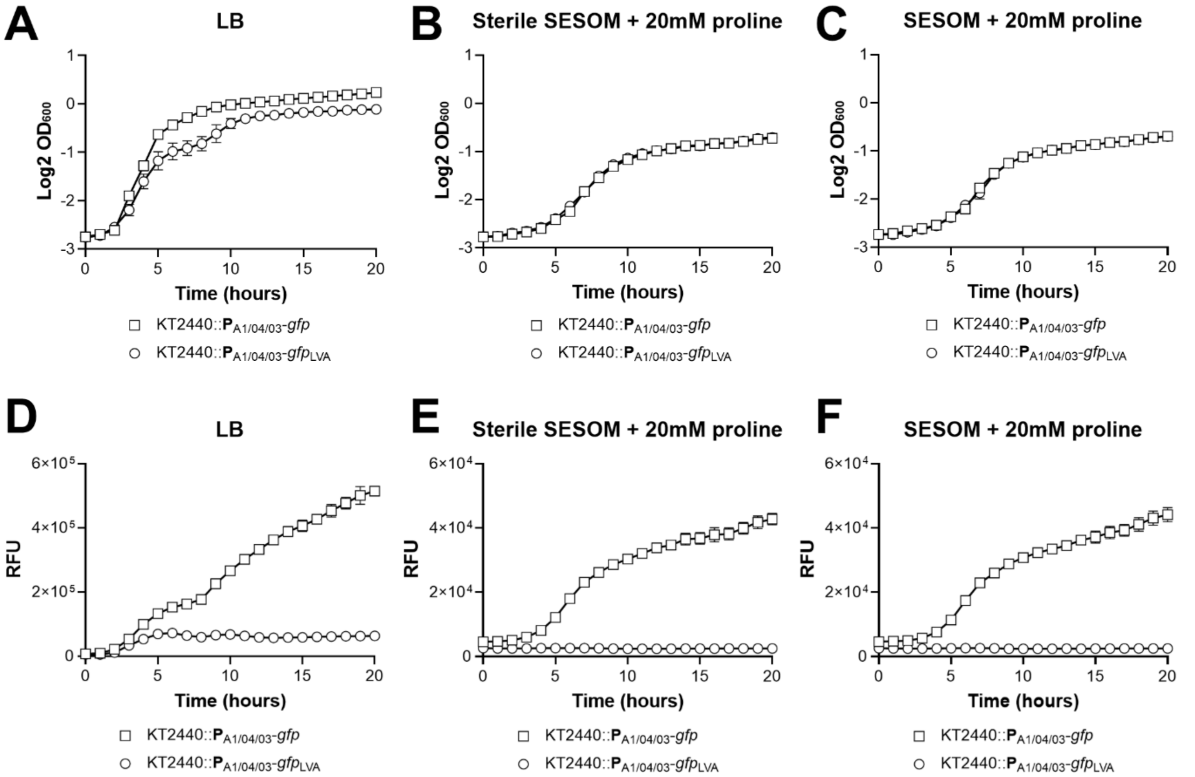


**Figure S3 |Protease-dependent protein degradation is upregulated in SESOM compared to LB.** Top graphs display growth curves of *P. putida* KT2440::$\mathbf{P}_{A1/04/03}$-*gfp* and KT2440::$\mathbf{P}_{A1/04/03}$-*gfp*$_{LVA}$ in LB (**A**), sterile SESOM supplemented with proline (**B**), and SESOM supplemented with proline (**C**). Bottom graphs (**D**-**F**) display relative fluorescent units (RFU) over time of the two strains. Data points represent mean values with error bars displaying standard deviations.

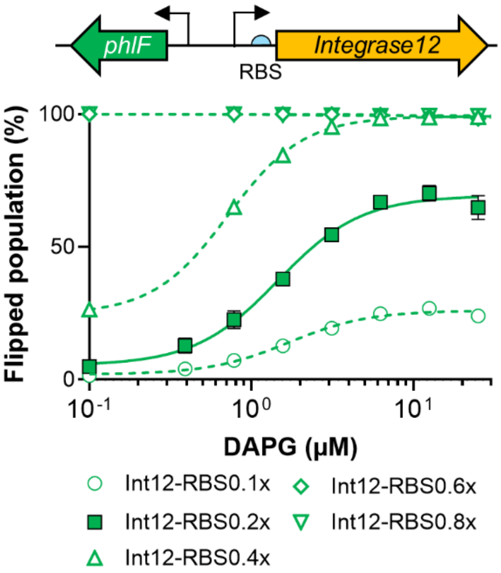


**Figure S4 | Removal of degradation tag and tuning Int12-RBS.** Removal of the protein degradation tag from *integrase12* and tuning its RBS instead. The DAPG system was used for tuning the RBS in LB with varying concentrations of DAPG. RBS0.2x (solid line) proved optimal with minimal leakiness in the absence of inducer, high dynamic range, and sensitivity. The left-most data points represent the recombination in the absence of inducer.

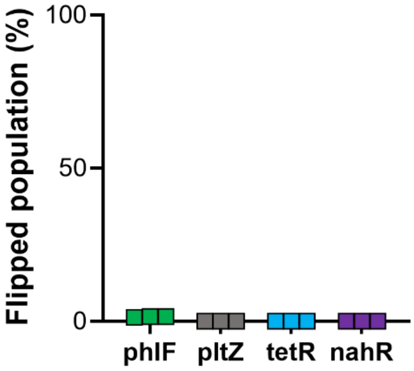


**Figure S5 | $T_0$ measurements of *P. putida* biosensors prior to SESOM assays.** Three biological replicates of each *P. putida* biosensor were cultivated in LB for 24 hours. After incubation, cultures were tested with flow cytometry to verify lack of leakiness in the absence of inducer prior to starting the SESOM assays.

Supplementary table 1 | Background strains

| Strain | Description | Source |
|---|---|---|
| *E. coli* TOP10 | Cloning | Invitrogen |
| *E. coli* DH10B | Assay strain | Invitrogen |
| *E. coli* CC118λpir | Cloning and conjugation | (1) |
| *E. coli* Pir1 | Conjugation for Tn7 transposon insertions | (2) |
| *P. putida* KT2440 | Wildtype | (3) |
| *P. putida* KT2440 Δall-Φ | Assay strain | (4) |
| *P. protegens* DTU9.1 | Wildtype | (5) |
| *P. protegens* DTU9.1 Δ*phlACB* | Deficient in DAPG production | (6) |
| *P. protegens* DTU9.1 Δ*pltA* | Deficient in pyoluteorin production | (6) |
| *P. protegens* DTU9.1 Δ*phlACB*, Δ*pltA* | Deficient in DAPG and pyoluteorin production | (6) |

Supplementary table 2 | Engineered strains

| Strain | Relevant genotype | Source |
|---|---|---|
| *P. putida* KT2440 | Tn5::**P**$_{A1/04/03}$-*gfp* | (7) |
| *P. putida* KT2440 | Tn5::**P**$_{A1/04/03}$-*gfp*$_{LVA}$ | (7) |
| *P. putida* KT2440 Δall-Φ | Rep12$_{PP_1035}$ | This study |
| *P. putida* KT2440 Δall-Φ | Rep12$_{PP_1035}$, Tn7::Int12$_{LVA}$-phlF | This study |
| *P. putida* KT2440 Δall-Φ | Rep12$_{PP_1035,}$ Tn7::Int12$_{RBS0.2x}$-phlF | This study |
| *P. putida* KT2440 Δall-Φ | Rep12$_{PP_1035,}$ Tn7::Int12$_{RBS0.2x}$-pltZ$_{RBS0.05x}$ | This study |
| *P. putida* KT2440 Δall-Φ | Rep12$_{PP_1035,}$ Tn7::Int12$_{RBS0.2x}$-tetR | This study |
| *P. putida* KT2440 Δall-Φ | Rep12$_{PP_1035,}$ Tn7::Int12$_{RBS0.2x}$-nahR | This study |
| *P. putida* KT2440 Δall-Φ | attB/P-DAPG barcode$_{PP_1035}$ | This study |
| *P. putida* KT2440 Δall-Φ | attL/R-**flipped**-DAPG barcode$_{PP_1035}$ | This study |
| *P. putida* KT2440 Δall-Φ | attB/P-Pyoluteorin barcode$_{PP_1035}$ | This study |
| *P. putida* KT2440 Δall-Φ | attB/P-aTc barcode$_{PP_1035}$ | This study |
| *P. putida* KT2440 Δall-Φ | attB/P-Salicylate barcode$_{PP_1035}$ | This study |
| *P. putida* KT2440 Δall-Φ | attB/P-DAPG barcode$_{PP_1035}$, Tn7::Int12$_{LVA}$-phlF | This study |
| *P. putida* KT2440 Δall-Φ | attB/P-Pyoluteorin barcode$_{PP_1035}$, Tn7::Int12$_{LVA}$-pltZ$_{RBS0.05x}$ | This study |
| *P. putida* KT2440 Δall-Φ | attB/P-aTc barcode$_{PP_1035}$, Tn7::Int12$_{LVA}$-tetR | This study |
| *P. putida* KT2440 Δall-Φ | attB/P-Salicylate barcode$_{PP_1035}$, Tn7::Int12$_{LVA,\ RBS4x}$-nahR | This study |

**Supplementary table 3** | Plasmids

| **Name** | **Source** |
|---|---|
| pRK600 | (8) |
| pTNS2 | (2) |
| pBG42-mKate | (5) |
| pRep12 (Original name: pReporter_12) | (9) |
| pAJM.847 | (10) |
| pAJM.011 | (10) |
| pAJM.771 | (10) |
| pInt12-araC (Original name: pIntegrase_12) | (9) |
| pInt12-phlF | This study |
| pInt12$_{ASV}$-phlF | This study |
| pInt12$_{LVA}$-phlF | This study |
| pSEVA231 | (11) |
| pSEVA231-Int12$_{LVA}$-phlF | This study |
| pSEVA231-Int12$_{LVA}$-pltZ$_{RBS0.05x}$ | This study |
| pSEVA231-Int12$_{LVA}$-tetR | This study |
| pSEVA231-Int12$_{LVA,\ RBS4x}$-nahR | This study |
| pSEVA231-Int12$_{LVA}$-fdeR$_{RBS0.1x}$ | This study |
| pSEVA231-Int12-araC | This study |
| pSEVA227-**P**$_{phlF}$-gfp | (12) |
| pTn7-M | (13) |
| pTn7- Int12$_{LVA}$-phlF | This study |
| pTn7- Int12$_{LVA}$-pltZ$_{RBS0.05x}$ | This study |
| pTn7- Int12$_{LVA}$-tetR | This study |
| pTn7- Int12$_{LVA,\ RBS4x}$-nahR | This study |
| pTn7- Int12$_{RBS0.2x}$-phlF | This study |
| pTn7- Int12$_{RBS0.2x}$-pltZ | This study |
| pTn7- Int12$_{RBS0.2x}$-tetR | This study |
| pTn7- Int12$_{RBS0.2x}$-nahR | This study |
| pK18msB | (14) |
| pMemoryArray | (9) |
| pK18msB-HR1/2$_{PP_1035}$-attB/P-DAPG barcode | This study |
| pK18msB-HR1/2$_{PP_1035}$-attL/R-**flipped**-DAPG barcode | This study |
| pK18msB-HR1/2$_{PP_1035}$-attB/P-Pyoluteorin barcode | This study |
| pK18msB-HR1/2$_{PP_1035}$-attB/P-aTc barcode | This study |
| pK18msB-HR1/2$_{PP_1035}$-attB/P-Salicylate barcode | This study |
| pK18msB$_{R6K}$ | This study |
| pK18msB$_{R6K}$-Rep12$_{PP_1035}$ | This study |

Supplementary table 4 | qPCR primers

| Primer name | Sequence |
| --- | --- |
| Prim1 | gttactaaaacaactggtcaagttc |
| Prim2 | gatctgagcgtaagttgaaatatg |
| Prim$_{DAPG}$ | gcactaggaagaattccgaag |
| Prim$_{Pyoluteorin}$ | ctatcgcgataattccatatcg |
| Prim$_{aTc}$ | atgaggctgcctgagatc |
| Prim$_{Salicylate}$ | acatctagttaatgtccgcaag |

52 Supplementary table 5 | Genetic parts

| Part name | Type | DNA sequence |
|---|---|---|
| **P**$_{J23119}$ | Promoter | ttgacagctagctcagtcctaggtataatgctagc |
| **P**$_{J23101}$ | Promoter | tttacagctagctcagtcctaggtattatgctagc |
| **P**$_{lacI}$ | Promoter | gcggcgcgccatcgaatggcgcaaaacctttcgcggtatggcatga<br>tagcgccc |
| **P**$_{lacIQ}$ | Promoter | gcggcgcgccatcgaatggtgcaaaacctttcgcggtatggcatga<br>tagcgccc |
| **P**$_{phlF}$ (10) | Promoter[a] | cgacgtacggtggaatctgattcgttaccaattgacatgatacgaa<br>acgtaccgtatcgttaaggt |
| **P**$_{pltZ}$ | Promoter[a] | gattcgttaccaattgacattaaattcaaattgaattttaattagg<br>cc |
| **P**$_{tetR}$ (10)<br>Original: **P**$_{Tet*}$ | Promoter[a] | ttttcagcaggacgcactgacctccctatcagtgatagagattgac<br>atccctatcagtgatagagatactgagcacag |
| **P**$_{nahR}$ (10)<br>Original: **P**$_{SalTTC}$ | Promoter[a] | gggcgcaatattcatgttgatgatttattatatatcgagtggtgta<br>tttatttatattgtttgctccgttaccgttattaac |
| **P**$_{fdeR}$ (15) | Promoter (upstream region) | gttgttgtgcttgttcttgccgaccctcggatagacgacggatggg<br>gtggtcaatgtattgatgccgtccatatcatgaatcaaaacaatcc<br>atttgatcaatatcaagctcactcttaagcttcactcatccgctgc<br>atggccccaccagaaagggctggcgcggcaagccggcggcgcactc<br>gcactggatgcgccgctgttgagcctggccatgacaacgcgccgat<br>agcggccacaccccgccaggcagggtaggagacaaggagacagac |
| **P**$_{14g}$ (13) | Promoter | gcccattgacaaggctctcgcggccaggtataattgcacga |
| **P**$_{BAD}$ | Promoter | aagaaaccaattgtccatattgcatcagacattgccgtcactgcgt<br>cttttactggctcttctcgctaaccaaaccggtaaccccgcttatt<br>aaaagcattctgtaacaaagcgggaccaaagccatgacaaaaacgc<br>gtaacaaaagtgtctataatcacggcagaaaagtccacattgatta<br>tttgcacggcgtcacactttgctatgccatagcattttttatccata<br>agattagcggatcctacctgacgctttttatcgcaactctctactg<br>tttctccat |
| **P**$_{A1/04/03}$ (7) | Promoter[a] | aaaatttatcaaaaagagtgttgacttgtgagcggataacaatgat<br>acttagattcaattgtgagcggataacaatttcacacatctagaat<br>t |
| RiboJ (16) | Ribozyme insulator | ttaaacaaaattatttgtagaggctgtttcgtcctcacggactcat<br>cagaccggaaagcacatccggtgacagct |
| BCD2 (17) | Bicistronic linker | gcccaagttcacttaaaaaggagatcaacaatgaaagcaattttcg<br>tactgaaacatcttaatcatgctaaggaggt |
| phl2 (10) | RBS[b] | ggaagagagtcaattcatgggggtgaat**atg** |
| BBa_B0030 | RBS[b] | aaagaggagaaatactag**atg** |
| BBa_B0034 | RBS[b] | aaagaggagaaatactag**atg** |
| tet1 (10) | RBS[b] | ggaagagagtcaattcagggtggtgaat**atg** |
| nah2 (10) | RBS[b] | ggaagagagtcaattcatggggttgaat**atg** |
| PltZ-RBS-0.05x | RBS[b] | tactagagaaagagcttgagtactag**atg** |
| PltZ-RBS-0.1x | RBS[b] | tactagagaaagagtccgactactag**atg** |

| | | |
|---|---|---|
| PltZ-RBS-0.4x | RBS[b] | tactagagaaagaggagccgtactag**atg** |
| PltZ-RBS-0.6x | RBS[b] | tactagagaaagagggagcctactag**atg** |
| FdeR-RBS-0.1x | RBS[b] | tactagagaaagaggttttgtactag**atg** |
| FdeR-RBS-0.25x | RBS[b] | tactagagaaagaggttgtatactag**atg** |
| FdeR-RBS-0.5x | RBS[b] | tactagagaaagaggggggcctactag**atg** |
| Int12-RBS (9) | RBS[b] | acacaggaagaaggctcg**atg** |
| Int12-RBS$_{nahR}$-4x | RBS[b] | atagcagaagaaggctcg**atg** |
| Int12-RBS$_{nahR}$-8x | RBS[b] | acgagagaagaaggctcg**atg** |
| Int12-RBS$_{nahR}$-16x | RBS[b] | aagggagaagaaggctcg**atg** |
| Int12-RBS-0.1x | RBS[b] | acacagggtacgggctcg**atg** |
| Int12-RBS-0.2x | RBS[b] | acacagtggcatggctcg**atg** |
| Int12-RBS-0.4x | RBS[b] | acacagagtagcggctcg**atg** |
| Int12-RBS-0.6x | RBS[b] | acacagtagtcgggctcg**atg** |
| Int12-RBS-0.8x | RBS[b] | acacaggatgggggctcg**atg** |
| ASV-tag (7) | Degradation tag | ggatccgcagcaaacgacgaaaactacgctgcatcagtt |
| LVA-tag (7) | Degradation tag | ggatccgcagcaaacgacgaaaactacgctttagtagct |
| *phlF* (10)<br>Original: *phlF*[AM] | Gene | atggcacgtaccccgagccgtagcagcattggtagcctgcgtagtc<br>cgcatacccataaagcaattctgaccagcaccattgaaatcctgaa<br>agaatgtggttatagcggtctgagcattgaaagcgtggcacgtcgc<br>gccggtgcaggcaaaccgaccatttatcgttggtggaccaacaaag<br>cagcactgattgccgaagtgtatgaaaatgaaatcgaacaggtacg<br>taaatttccggatttgggtagctttaaagccgatctggattttctg<br>ctgcataatctgtggaaagtttggcgtgaaaccatttgtggtgaag<br>catttcgttgtgttattgcagaagcacagttggaccctgtaaccct<br>gacccaactgaaagatcagtttatggaacgtcgtcgtgagataccg<br>aaaaaactggttgaagatgccattagcaatggtgaactgccgaaag<br>atatcaatcgtgaactgctgctggatatgatttttggtttttgttg<br>gtatcgcctgctgaccgaacagttgaccgttgaacaggatattgaa<br>gaatttaccttcctgctgattaatggtgtttgtccgggtacacagt<br>gttga |
| *pltZ* | Gene | atgaagcaaccaccagctcagactacgcgtaatattatgaatactc<br>gccgccgtacaagccgttctgacggcgagcacacaaaaattcgtat<br>tttagaggtagcggcacgtttgtttgcgcaacacgggtatgcaaat<br>actgcatctaagttaatctgtgaggaagctggggccgacctggcgg<br>cgattaattatcatttcggaagtcgtgaggccttatacaaagctgt<br>attgattgagggccataaacaactggtatcttttgaagccttgagc<br>caattagcgcaatctgaagagccagcccttatcaagctggacagct<br>ttattgacgcgattgtaacacgtgtacttgacgagcaaagttggca<br>gagcaaagtttgcgcgcgtgagattttggctcccactgtgcatttt<br>acgagcttggtccaggaggaagttatgccgaaatttcgtctgttag<br>aagctctgatctccgagatcactggctttccaattggtgatccggc |

| | | |
|---|---|---|
| | | cttggcccgttgcactatctccattattgcaccctgcttaatgtta<br>gctgtaattgatcgtcagcaacctagtccattgcaagccgttttac<br>agcatgatgctaatgctctgaaagcacactttaagttatttgctcg<br>ttcgggtttggcagcaatcgcgcagtag |
| *tetR* (10) | Gene | atgtccagattagataaaagtaaagtgattaacagcgcattagagc<br>tgcttaatgaggtcggaatcgaaggtttaacaacccgtaaactcgc<br>ccagaagctaggtgtagagcagcctacattgtattggcatgtaaaa<br>aataagcgggctttgctcgacgccttagccattgagatgttagata<br>ggcaccatactcacttttgccctttagaaggggaaagctggcaaga<br>ttttttacgtaataacgctaaaagttttagatgtgctttactaagt<br>catcgcgatggagcaaaagtacatttaggtacacggcctacagaaa<br>aacagtatgaaactctcgaaaatcaattagcctttttatgccaaca<br>aggtttttcactagagaatgcattatatgcactcagcgctgtgggg<br>cattttactttaggttgcgtattggaagatcaagagcatcaagtcg<br>ctaaagaagaaagggaaacacctactactgatagtatgccgccatt<br>attacgacaagctatcgaattatttgatcaccaaggtgcagagcca<br>gccttcttattcggccttgaattgatcatatgcggattagaaaaac<br>aacttaaatgtgaaagtgggtcctga |
| *nahR* (10)<br>Original: *nahR*$^{AM}$ | Gene | atggaactgcgtgacctggatttaaacctgctggtggtgttcaacc<br>agttgctggtcgacagacgcgtctctgtcactgcggagaacctggg<br>cctgacccagcctgccgtgagcaatgcgctgaaacgcctgcgcacc<br>tcgctacaggacccactcttcgtgcgcacacatcagggaatggaac<br>ccacaccctatgccgcgcatctggccgagcacgtcacttcggccat<br>gcacgcactgcgcaacgccctacagcaccatgaaagcttcgatccg<br>ctgaccagcgagcgtaccttcaccctggccatgaccgacattggcg<br>agatctacttcatgccgcggctgatggatgcgctggctcaccaggc<br>ccccaattgcgtgatcagtacggtgcgcgacagttcgatgagcctg<br>atgcaggccttgcagaacggaaccgtggacttggccgtgggcctgc<br>ttcccaatctgcaaactggcttctttcagcgccggctgctccagaa<br>tcactacgtgtgcctatgtcgcaaggaccatccagtcacccgcgaa<br>cccctgactctggagcgcttctgttcctacggccacgtgcgtgtca<br>tcgccgctggcaccggccacggcgaggtggacacgtacatgacacg<br>ggtcggcatccggcgcgacatccgtctggaagtgccgcacttcgcc<br>gccgttggccacatcctccagcgcaccgatctgctcgccactgtgc<br>cgatatgtttagccgactgctgcgtagagcccttcggcctaagcgc<br>cttgccgcacccagtcgtcttgcctgaaatagccatcaacatgttc<br>tggcatgcgaagtaccacaaggacctagccaatatttggttgcggc<br>aactgatgtttgacctgtttacggattga |
| *fdeR* | Gene | atgcgcttcaataaattagatctgaacttgcttgtggcgcttgatg<br>cattgctgacggagatgtctatctcacgcgctgccgaaaaaattca<br>tctgtcacaatctgccatgtcgaacgccttggcgcgcttgcgtgaa<br>tacttcgatgatgaattgttaatccaggtaggacgccgcatggaac<br>ctacaccccgcgcagaggtcctgaaggatgccgtgcacgatgtctt<br>acgccgtatcgatggtagtattgcggctcttccagcgttcgtcccg<br>gctgagtcaacccgcgagtttcgcatctcggtcagtgatttcactt<br>tatctgtcttgatcccccgcgttttggctcgtgcacacgcagaagg<br>aaaacatattcgctttgcactgatgccacaagtgcaggacccaacc<br>cgttcactggaccgtgccgaagtggatttgttagtcttaccgcaag<br>agttctgtactcctgaccaccctgccgaggaggtgttccgtgagcg<br>ccacgtgtgtgtagtgtggcgcgatagcgctctggctcaaggggag<br>ttgacgttggaacgctatatggcgtcaggccatgtagtaatggttc<br>cacctggagccaatgcaagttcggtcgaagcatggatggcacgcaa<br>attaggtttcgcgcgtcgcgtggaagtgacttctttttcctttgca<br>tcggcgcttgccttggtgcaagggaccgatcgcattgcgactgttc<br>atgcacgtcttgcacagcttctggcacctcagtggccggttgtaat<br>taaagagtctcctttgtctttaggggaaatgcgccaaatgatgcag<br>tggcaccgttatcgctcgaatgatccgggtattcagtggttacgtc<br>gtgtctttcttgagtcagcccaggagatggatgcggctcttccggg<br>catttgctga |
| *araC* (9) | Gene | atggctgaagcgcaaaatgatcccctgctgccgggatactcgttta<br>atgcccatctggtggcgggtttaacgccgattgaggccaacggtta |

| | | |
|---|---|---|
| | | tctcgatttttttatcgaccgaccgctgggaatgaaaggttatatt<br>ctcaatctcaccattcgcggtcagggggtggtgaaaaatcagggac<br>gagaatttgtttgccgaccgggtgatattttgctgttcccgccagg<br>agagattcatcactacggtcgtcatccggaggctcgcgaatggtat<br>caccagtgggtttactttcgtccgcgcgcctactggcatgaatggc<br>ttaactggccgtcaatatttgccaatacggggttctttcgcccgga<br>tgaagcgcaccagccgcatttcagcgacctgtttgggcaaatcatt<br>aacgccgggcaaggggaagggcgctattcggagctgctggcgataa<br>atctgcttgagcaattgttactgcggcgcatggaagcgattaacga<br>gtcgctccatccaccgatggataatcgggtacgcgaggcttgtcag<br>tacatcagcgatcacctggcagacagcaattttgatatcgccagcg<br>tcgcacagcatgtttgcttgtcgccgtcgcgtctgtcacatctttt<br>ccgccagcagttagggattagcgtcttaagctggcgcgaggaccaa<br>cgtatcagccaggcgaagctgcttttgagcaccacccggatgccta<br>tcgccaccgtcggtcgcaatgttggttttgacgatcaactctattt<br>ctcgcgggtatttaaaaaatgcaccggggccagcccgagcgagttc<br>cgtgccggttgtgaagaaaaagtgaatgatgtagccgtcaagttgt<br>cataa |
| *integrase12* (9) | Gene | atgaaagtggccatttatacccgtgttagcagcgcagaacaggcaa<br>atgaaggttatagcattcacgagcagaagaagaaactgatcagcta<br>ttgcgaaatccacgattggaacgagtataaagtttttaccgatgca<br>ggtattagcggtggtagcatgaaacgtccggcactgcaaaaactga<br>tgaaacatctgagttcatttgatctggtgctggtgtataaactgga<br>tcgtctgacccgtaatgttcgtgatctgctggatatgctggaagaa<br>tttgaacagtataacgtgagctttaaaagcgccaccgaagtttttg<br>ataccaccagtgcaattggcaaactgtttattaccatggttggtgc<br>aatggcagaatgggaacgtgaaaccattcgtgaacgtagcctgttt<br>ggtagccgtgcagcagttcgtgaaggtaactatattcgtgaagcac<br>cgttttgctatgataacattgaaggtaaactgcacccgaacgaata<br>tgccaaagttattgatctgattgtgagcatgttcaaaaaaggcatt<br>agcgccaatgaaattgcacgtcgtctgaatagcagcaaagttcatg<br>ttccgaacaaaaaaagctggaatcgtaatagcctgattcgtctgat<br>gcgtagtccggttctgcgtggtcataccaaatatggtgatatgctg<br>attgaaaacacccatgaaccggtgctgagcgaacatgattataatg<br>caattaacaacgccatcagcagcaaaacccataaaagcaaagttaa<br>acaccatgccatttttcgtggtgcactggtttgtccgcagtgtaat<br>cgtcgtctgcatctgtatgcaggcaccgttaaagatcgtaaaggct<br>ataaatacgatgtgcgtcgctataaatgtgaaacctgcagcaaaaa<br>caaagatgtgaagaatgtgagcttcaacgaaagcgaagtggaaaac<br>aaattcgtcaatctgctgaaaagctacgagctgaacaaatttcata<br>tccgtaaagtggaaccggtgaaaaaaatcgagtatgacatcgataa<br>gattaacaaacagaaaattaactatacccgcagttggagcctgggc<br>tatattgaagatgatgaatatttcgagctgatggaagaaatcaacg<br>ccaccaaaaaaatgatcgaagaacagaccaccgagaataaacagag<br>cgttagcaaagagcagattcagagcattaacaactttatcctgaaa<br>ggctgggaagaactgaccatcaaagataaagaggaactgattctga<br>gcaccgtggataaaatcgaatttaacttcatcccgaaagataaaaa<br>acataaaaccaataccctggatattaacaatattcactttaaattc<br>taa |
| *gfp* (9) | Gene | atgagtaaaggagaagaacttttcactggagttgtcccaattcttg<br>ttgaattagatggtaatgttaatgggcacaaattttctgtcagtgg<br>agagggtgaaggtgatgcagcatacggaaaacttacccttaaattt<br>atttgcactactggaaaactacctgttccatggccaacacttgtca<br>ctactttgacttatggtgttcaatgcttttcaagatacccagatca<br>tatgaaacggcatgactttttcaagagtgccatgcccgaaggttat<br>gtacaggaaagaactatatttttcaaagatgacgggaactataaga<br>cacgtgctgaagtcaagtttgaaggtgatacacttgttaatagaat<br>cgagttaaaaggtattgattttaaagaagatggaaacattcttgga<br>cacaagttggaatacaactataactcacacaatgtatacatcatgg<br>cagacaaacaaaagaatggaatcaaagttaacttcaaaattagaca<br>caacattgaagatggaagcgttcaactagcagaccattatcaacaa |

| | | |
|---|---|---|
| | | aatactccaattggcgatggccctgtccttttaccagacaaccatt<br>acctgtccacacaatctgccctttcgaaagatcccaacgaaaagag<br>agaccacatggtccttcttgagtttgtaacagctgctgggattaca<br>catggcatggatgaactatacaaa |
| *dsRed* (9) | Gene | atggcttcctccgaagacgttatcaaagagttcatgcgtttcaaag<br>ttcgtatggaaggttccgttaacggtcacgagttcgaaatcgaagg<br>tgaaggtgaaggtcgtccgtacgaaggtacccagaccgctaaactg<br>aaagttaccaaaggtggtccgctgccgttcgcttgggacatcctgt<br>ccccgcagttccagtacggttccaaagcttacgttaaacacccggc<br>tgacatcccggactacctgaaactgtccttcccggaaggtttcaaa<br>tgggaacgtgttatgaacttcgaagacggtggtgttgttaccgtta<br>cccaggactcctccctgcaagacggtgagttcatctacaaagttaa<br>actgcgtggtaccaacttcccgtccgacggtccggttatgcagaaa<br>aaaaccatgggttgggaagcttccaccgaacgtatgtacccggaag<br>acggtgctctgaaaggtgaaatcaaaatgcgtctgaaactgaaaga<br>cggtggtcactacgacgctgaagttaaaaccacctacatggctaaa<br>aaaccggttcagctgccgggtgcttacaaaaccgacatcaaactgg<br>acatcacctcccacaacgaagactacaccatcgttgaacagtacga<br>acgtgctgaaggtcgtcactccaccggtgcttaa |
| *mKate* (18) | Gene | atgatcatgtcagaacttatcaaggaaaatatgcacatgaaattgt<br>acatggaaggaacagtaaataatcaccactttaaatgtacctcaga<br>aggagaaggaaaaccatatgaaggtactcaaaccatgcgtattaag<br>gccgttgaaggtggaccattgccttttgcctttgatattcttgcta<br>catcttttatgtacggatcaaagacttttatcaatcatacccaagg<br>tatcccagatttctttaaacagtcatttcctgaaggatttacatgg<br>gaacgtgtcacaacttatgaagatggtggagtattgacagcaactc<br>aagatacatctttgcaagatggttgtcttatctacaatgtaaagat<br>ccgtggagttaattttccaagtaatggtcctgttatgcagaaaaag<br>acccttggatgggaagcatctaccgaaacattatatcctgccgatg<br>gtggattggaaggtcgtgctgatatggcattgaaacttgtcggtgg<br>aggtcaccttatctgtaatttgaagaccacataccgttctaaaaag<br>ccagctaagaatcttaagatgcctggtgtttactacgtcgatcgtc<br>gtttagaacgtatcaAagaagcagataaggaaacttatgttgaaca<br>gcacgaagtagccgtcgcacgttattgtgatttgcctagtaaattg<br>ggacaccgttaa |
| *attB*$_{Int12}$ (9) | Integrase recognition site | gttcgtggtaactatgggtggtacaggtgccacattagttgtacca<br>tttatgtttatgtggttaac |
| *attP*$_{Int12}$ (9) | Integrase recognition site | ttaaattttaatagcagttgtgtcactatttaggtctacccagtga<br>cacaactaacatacaaaa |
| attB/P-DAPG barcode<br>Original: Spacer 12 | DNA barcode[c] | gttcgtggtaactatgggtggtacaggtgccacattagttgtacca<br>tttatgtttatgtggttaacgggaccttcggaattcttcctagtgc<br>gcgaccgacggagtgttcctctggttaaattttaatagcagttgtg<br>tcactatttaggtctacccagtgacacaactaacatacaaaa |
| attL/R-**flipped**-DAPG barcode | DNA barcode[c] | gttcgtggtaactatgggtggtacaggtgccacctaaatagtgaca<br>caactgctattaaaatttaagggaccttcggaattcttcctagtgc<br>gcgaccgacggagtgttcctctgggttaaccacataaacataaatg<br>gtacaactaatgtctacccagtgacacaactaacatacaaaa |
| attB/P-Pyoluteorin barcode<br>Original: Spacer 4 | DNA barcode[c] | gttcgtggtaactatgggtggtacaggtgccacattagttgtacca<br>tttatgtttatgtggttaactgtacgatatggaattatcgcgatag<br>gatacgacctctaatgttcacaacttaaattttaatagcagttgtg<br>tcactatttaggtctacccagtgacacaactaacatacaaaa |
| attB/P-aTc barcode<br>Original: Spacer 9 | DNA barcode[c] | gttcgtggtaactatgggtggtacaggtgccacattagttgtacca<br>tttatgtttatgtggttaacttctagaggatctcaggcagcctcat<br>gctgatcgccaaggcaatggtgctttaaattttaatagcagttgtg<br>tcactatttaggtctacccagtgacacaactaacatacaaaa |
| attB/P-Salicylate barcode<br>Original: Spacer 5 | DNA barcode[c] | gttcgtggtaactatgggtggtacaggtgccacattagttgtacca<br>tttatgtttatgtggttaacgagtcttgcggacattaactagatgt<br>gcgcagcgggcatcaaatctttttttaaattttaatagcagttgtg<br>tcactatttaggtctacccagtgacacaactaacatacaaaa |

| | | |
|---|---|---|
| $HR1_{PP\_1035}$ | Homology region | ctccaaccgtcacgacatcaaggaaccgtggttcaccgacacctac<br>ttgctgggcaagaacgagcgcgcccaggtagcactggtggcggata<br>accctggaacctggatgtttcactgccacgtcatcgaccacatgga<br>aaccggcctgatggccgcgattgcggtggtctgatgcgcccgcaga<br>tcatcgatcgcagtcgcgatcaggaattcatgcgcctggcgctggc<br>cttggccgccgaaggcgcggcgttgggcgaggtgccagtgggggcg<br>gtgctggtgcagcatgggcaggtgatcggccagggtttcaaccggc<br>cgatcatcgacagcgaccccagcgcgcatgccgagatggtggcgat<br>cagggcggcagccaggtcggccagcaactaccggctacccggcagc<br>accctgtatgtgaccctggaaccgtgcagcatgtgtgcggggctga<br>tcgtgcattcgcgggtgatgcgcgtggtgtttggcgcgctggagcc<br>caaggccgggatcgtgcagagccaggggcagttctttggccaaggc<br>tttctcaaccaccgggtgatggtggagggcggggtgctggcgcaag<br>agtgcgggcagatcctcagcgacttcttcaaggcccgccgggccaa<br>ggcttgagctgcaccggcctcttcgcgggcaagcccgctcctacag<br>gttctgcgcaaatcttgtaggcgcgggtt |
| $HR2_{PP\_1035}$ | Homology region | tgcccgcgaagaggtcctgacaggcttacatcggcgacgacttatc<br>cgctggcttgtccttgttgaccccgggcacatgcaggttgccctcg<br>gccacctggccttccagctgcggctgggtcacccaggtgaggatgt<br>cgtagtaacggcggatgttggccacgaaatgcaccggttcgccccc<br>acgggcatagccgtacttggtctgccggtaccactgcttctgagcc<br>agccgcggcagcatcttcttcacgtccagccacttgttcgggttca<br>gcttctcgcgcttggccagggtgcgggcgtcttccaggtggccgcc<br>accgacattgtaggcggcaagggcgaaccaggtgcggtccggctcc<br>ttgatgctgtcgtcaagctcttccttgatcttcatgaagtacttgg<br>cgccgccctggatgctctgcctcgggtccagtcggttcgacacgcc<br>catggcctgggcggtgcgctgggtcagcatcatcaggccgcgcacg<br>ccggtcttggaggtgacttccggctgccacatcgattcctggtagc<br>caatggctgccagcaggcgccagtcgacctgctcgaccttggcgta<br>gctcttgaagtgcttctcgtacttgggcaggcgctgttgcaggtgc<br>tgggcgaaggtgtaggcgccgacatagccaagtacgtcgacatggc<br>cgtagtagcggtccttcaatcgctgaagggtgccgtttttctgtgc<br>ctt |

[a] Red color indicates the known operator site for regulator binding

[b] Bold indicates start codon downstream of RBS sequences

[c] Underlines indicates the unique 50bp barcode for each sensor
